# Selective and mechanistic sources of recurrent rearrangements across the cancer genome

**DOI:** 10.1101/187609

**Authors:** Jeremiah A Wala, Ofer Shapira, Yilong Li, David Craft, Steven E Schumacher, Marcin Imielinski, James E Haber, Nicola D Roberts, Xiaotong Yao, Chip Stewart, Cheng-Zhong Zhang, Jose Tubio, Young Seok Ju, Peter J Campbell, Joachim Weischenfeldt, Rameen Beroukhim, on behalf of the PCAWG-Structural Variation Working Group and the PCAWG Network.

**Affiliations:** Broad Institute, Cambridge, MA 02142, USA; Department of Cancer Biology, Dana-Farber Cancer Institute, Boston, MA 02215, USA; Bioinformatics and Integrative Genomics, Harvard University, Cambridge, MA, USA; SBGD Inc., Cambridge, MA 02142, USA; Cancer Genome Project, Wellcome Trust Sanger Institute, Wellcome Trust Genome Campus, Hinxton, Cambridgeshire CB10 1SA UK; Department of Radiation Oncology, Harvard Medical School, Massachusetts General Hospital, 55 Fruit St., Boston, MA 02114, USA; New York Genome Center, New York, NY 10013, USA; Department of Pathology and Laboratory Medicine, Englander Institute for Precision Medicine, Institute for Computational Biomedicine, and Meyer Cancer Center, Weill Cornell Medicine, New York, NY 10065, USA; Department of Biology and Rosenstiel Basic Medical Sciences Research Center, Brandeis University, Waltham, MA 02454; Department of Biostatistics and Computational Biology, Dana-Farber Cancer Institute, 44 Binney Street DA1410, Boston, MA 02115, USA; Department of Biomedical Informatics, Harvard Medical School, 10 Shattuck Street 315B, Boston, MA 02115, USA; Mobile Genomes & Disease, The Biomedical Research Centre - CINBIO, University of Vigo, 36310 Vigo, Spain; Department of Haematology, University of Cambridge, Cambridge CB2 2XY, UK; Biotech Research & Innovation Centre (BRIC); The Finsen Laboratory, Rigshospitalet, University of Copenhagen, Copenhagen, Denmark; European Molecular Biology Laboratory (EMBL), Genome Biology Unit, Heidelberg, Germany; Department of Medical Oncology, Dana-Farber Cancer Institute, Boston, MA 02215, USA

**Author notes:** These authors contributed equally to this work (Jeremiah A Wala, Ofer Shapira). **Co-corresponding authors**: Peter J Campbell, MD, PhD; Cancer Genome Project, Wellcome Trust Sanger Institute; Hinxton, Cambridgeshire CB10 1SA, United Kingdom, Joachim Weischenfeldt, PhD; Biotech Research & Innovation Centre (BRIC); The Finsen Laboratory, Rigshospitalet, University of Copenhagen, Copenhagen, Denmark, Rameen Beroukhim, MD, PhD; Dana-Farber Cancer Institute; 450 Brookline Avenue, Smith 1022C, Boston, MA, 02115, USA.

**Keywords:** structural variation, rearrangements, cancer genomics, pan-cancer

## Abstract

Cancer cells can acquire profound alterations to the structure of their genomes, including rearrangements that fuse distant DNA breakpoints. We analyze the distribution of somatic rearrangements across the cancer genome, using whole-genome sequencing data from 2,693 tumor-normal pairs. We observe substantial variation in the density of rearrangement breakpoints, with enrichment in open chromatin and sites with high densities of repetitive elements. After accounting for these patterns, we identify significantly recurrent breakpoints (SRBs) at 52 loci, including novel SRBs near *BRD4* and *AKR1C3*. Taking into account both loci fused by a rearrangement, we observe different signatures resembling either single breaks followed by strand invasion or two separate breaks that become joined. Accounting for these signatures, we identify 90 pairs of loci that are significantly recurrently juxtaposed (SRJs). SRJs are primarily tumor-type specific and tend to involve genes with tissue-specific expression. SRJs were frequently associated with disruption of topology-associated domains, juxtaposition of enhancer elements, and increased expression of neighboring genes. Lastly, we find that the power to detect SRJs decreases for short rearrangements, and that reliable detection of all driver SRJs will require whole-genome sequencing data from an order of magnitude more cancer samples than currently available.

## Introduction

Rearrangements can substantially alter the structure and function of the genome and underlie many strongly oncogenic driver alterations in cancer^1^. A single rearrangement involves both the breaking of DNA and the formation of novel aberrant juxtapositions between pairs of genomically distant breakpoints, often affecting large stretches of DNA in a single event. Rearrangement topologies may be relatively simple (deletions, inversions, duplications or balanced translocations), or highly complex, as in the chromosomal shattering of chromothripsis^2^ or clustered reciprocal rearrangements of chromoplexy^3^. A single rearrangement can generate novel fusion gene products, affect gene expression by altering gene copy number (dosage effects), or affect gene expression by altering gene regulatory networks in *cis*^*4*^. Cis-regulatory effects of a rearrangement can alter expression of genes up to two megabases from the event locus^5^, and the changes in copy-number induced by a single rearrangement can span over one hundred megabases. A complete understanding of the landscape of selective pressures for rearrangements must therefore account for both loss-of-function and gain-of-function effects, and the possibility that a single rearrangement can alter two or more genes simultaneously^6^.

Whole-genome sequencing data are required to detect rearrangements genome-wide, as the vast majority of rearrangement breakpoints lie outside of exons. As such, large pan-cancer rearrangement analyses have been limited by the relatively small numbers of cancers profiled by whole-genome sequencing. However, several questions are often more easily addressed in pan-cancer analyses, due to the size of the datasets involved and the ability to compare data from different cancer types (https://doi.org/10.1101/162784).

Here we assess rearrangements across 2,693 cancer whole-genomes from 30 histological subtypes as part of the Pan-Cancer Analysis of Whole Genomes (PCAWG). We identify genome-wide patterns that both predict the distribution of breakpoints and rearrangements and inform the mechanisms by which they are formed. Accounting for these patterns, we discover significantly recurrent breakpoints (SRBs) and significantly recurrent juxtapositions (SRJs) between pairs of loci brought together by a rearrangement. These recurrent events include both known and novel candidate drivers and are associated with substantial changes in gene expression. We further find that the recurrent SRJs, more than other types of somatic genetic events, are strongly associated with cell-of-origin, suggesting that SRJs are shaped by the epigenetic state of the cell. Finally, we calculate the statistical power required to identify SRJs. We show that this is highly dependent on the genomic distance between the two breakpoints of a rearrangement, and find that, when taking into account all distances, we are substantially underpowered to detect important events.

## Results

### Rearrangement density along the genome is determined most strongly by chromatin structure and sequence features

We analyzed 292,253 high-confidence somatic rearrangements (584,506 breakpoints) in 2,693 cancer-whole genomes across 30 histological cancer subtypes as part of the Pan-Cancer Analysis of Whole Genomes (PCAWG) of the International Cancer Genome Consortium (ICGC) (**Supp. Note; Supp. Figs. 1-3**). The effects of each of these rearrangements may derive primarily from the disruption of genomic locations of one or both of their *breakpoints* (such as disruption of a tumor suppressor) or may result from the generation of a novel *juxtaposition* between loci due to reorganization of the genome, as in the case of *BRAF*-*KIAA1549* fusions. We therefore analyzed both where breakpoints tended to occur (the one-dimensional analysis) and which pairs of loci tended to be juxtaposed (the two-dimensional analysis; **Fig. 1a**).

**Figure 1:**
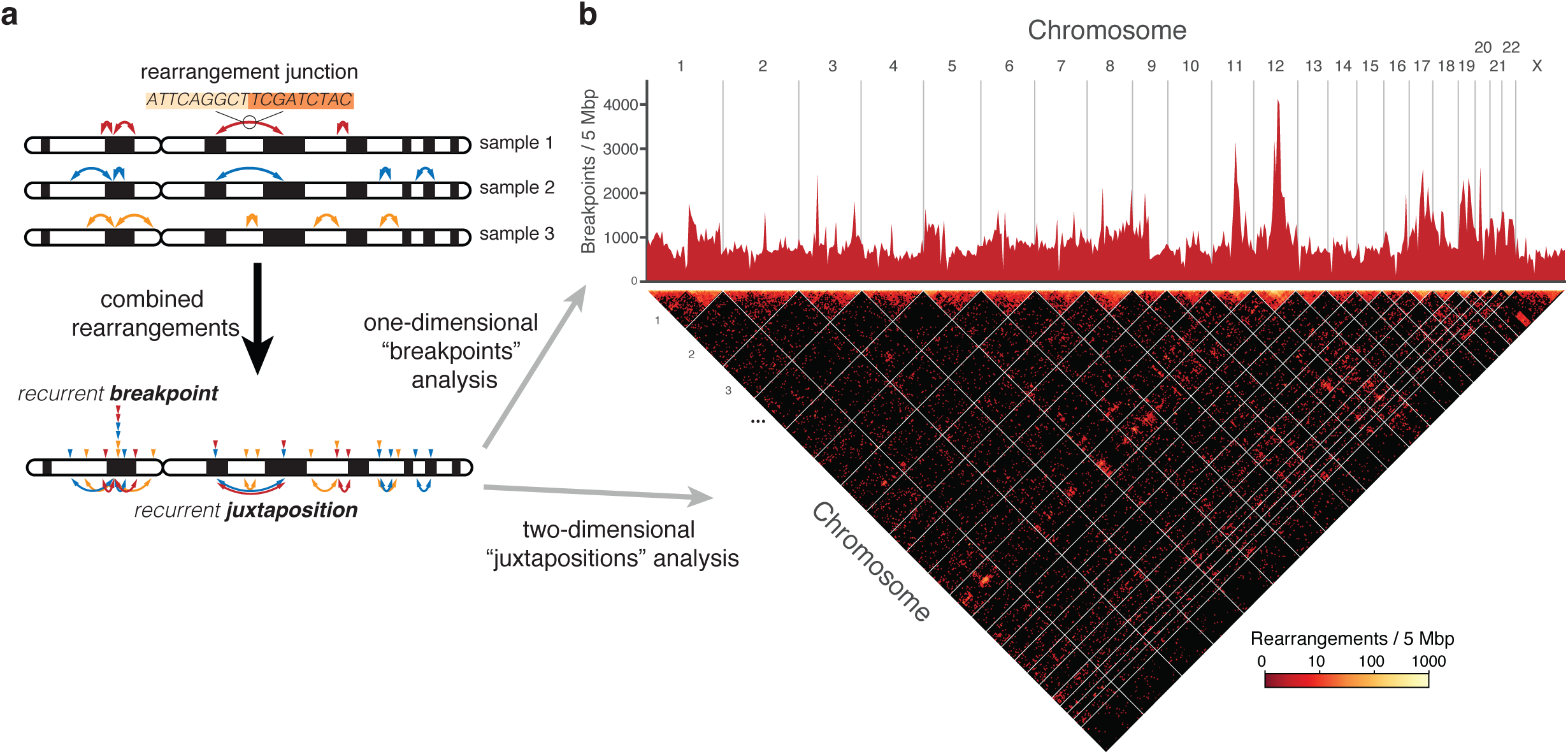
Analysis overview. a) Schematic indicating rearrangements and rearrangement junctions in three hypothetical genomes (top) and the two analysis approaches (bottom): the 1D analysis for recurrent breakpoints and the 2D analysis for recurrent juxtapositions between pairs of loci. b) The 1D density of breakpoints genome-wide (top) and 2D density of juxtapositions (bottom) across 2,693 cancer genomes.

Sources of variation in the 1D and 2D rearrangement densities could be from mechanistic biases towards rearrangements involving certain genomic loci, or from selective pressures that lead to enrichment of specific rearrangements among cancers. We first evaluated genome-wide patterns suggestive of mechanistic pressures and then identified specific loci where rearrangements were significantly enriched. This provided a catalog of candidate driver SRBs in the 1D analysis and SRJs in the 2D analysis events.

For the 1D analysis, we modeled genome-wide background variations in breakpoint density (**Fig. 1b**, top) using a gamma-Poisson (GP) model^7^ where DNA breaks follow a Poisson distribution that can vary based on local genomic features (see **Methods**; **Supp. Fig. 4, 5; Supp. Table 1**). We found that the density of short-interspersed nuclear elements (SINEs), fragile sites, gene expression, replication timing, and DNAase hypersensitivity sites were significant predictors of increased breakpoint density. Early replication timing, H3K36me3 density, GC content and heterochromatin were significantly predictive of decreased breakpoint densities (**Ext. Fig. 1**).

For the 2D analysis, we determined how the density of rearrangements between any two loci (**Fig. 1b**, bottom) varied with the genomic distance between those loci (the rearrangement’s “span”), the breakpoint density and sequence features of the two loci, and rearrangement topology.

Most rearrangements are short. Between 2 Kbp to 20 Mbp, the frequency of rearrangements is approximately inversely proportional to the rearrangement span (**Fig. 2a**), and the probability density drops by four orders of magnitude over this distance. This is similar to the global contact probability as a function of genomic distance determined by Hi-C mapping of non-cancer genomes^8^, and distributions of lengths for somatic copy-number alterations in cancer^9^,^10^. Rearrangements also tend to stay within the same topologically associated domain (TAD; **Fig. 2b**), supporting the role of three-dimensional chromatin organization in partner selection^9,11^.

**Figure 2:**
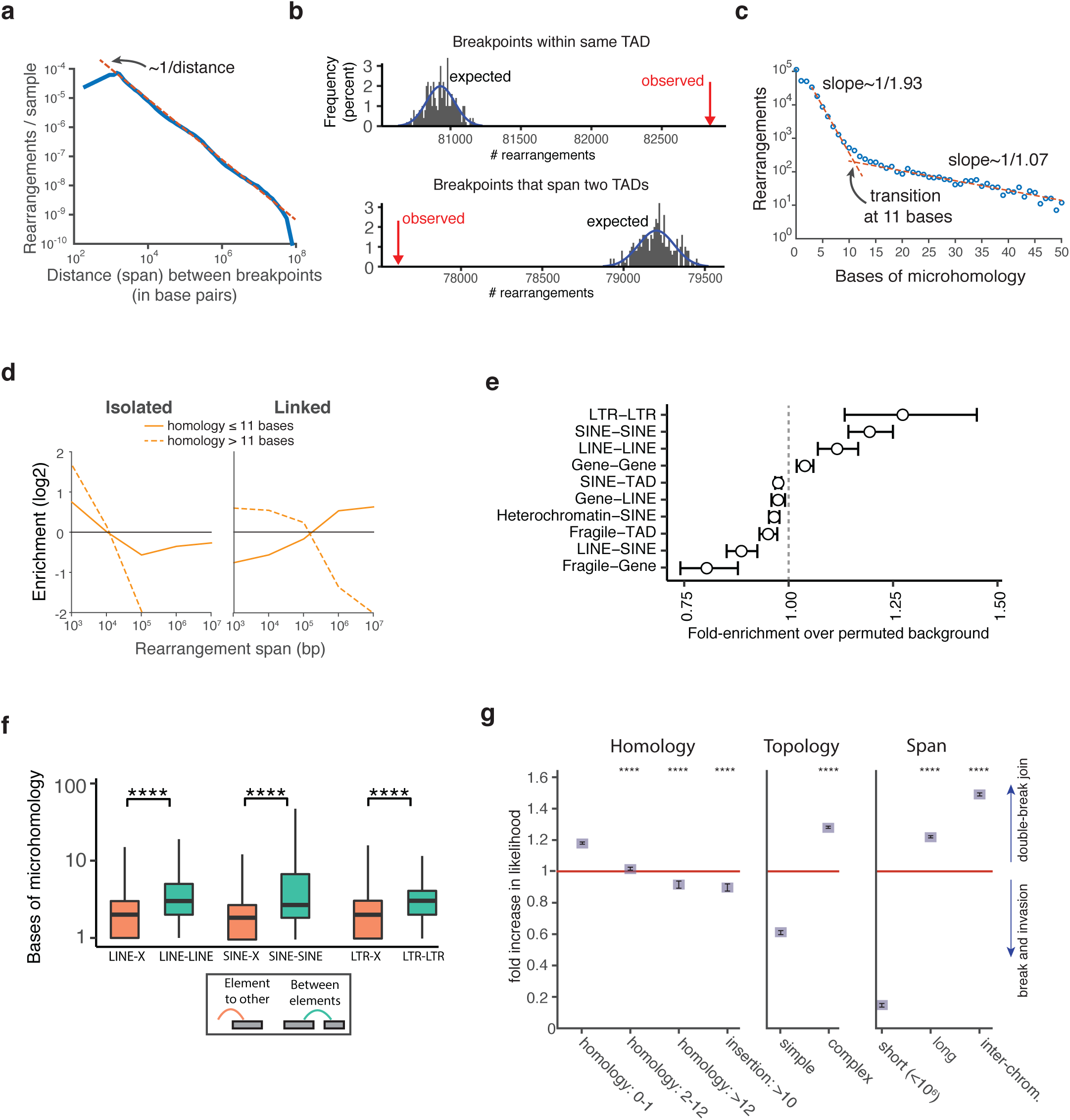
Determinants of the 2D density of rearrangements across the cancer genome. a) The distribution of spans (distances between breakpoint; x-axis) for intra-chromosomal rearrangements, superimposed upon an inverse power law distribution. b) Observed (red arrows) and expected distribution of (gray) numbers of rearrangements with both breakpoints within the same TAD (topologically associated domain; top)or with breakpoints that cross TADs (bottom). The expected distribution is based on permuted data. c) The frequency of rearrangements as a function of bases of microhomology. d) Enrichment of rearrangements categorized by topology (isolated, left; linked, right) and bases of microhomology relative to all rearrangements of similar span, as a function of rearrangement span (horizontal axis). e) Fold-enrichment or depletion for rearrangements between different elements for nine different genomic relationships, compared with the permuted background. Error bars represent 3 standard deviations of the fitted background distribution. f) Breakpoint microhomology for rearrangements connecting repetitive elements of the same class (green) or rearrangements with only one breakpoint in a repetitive element (orange). Comparisons with four stars indicate p<0.0001. g) Likelihood that rearrangements were generated by two breaks followed by a join, divided by the likelihood they were generated by a single break followed by strand invasion, for subsets of rearrangements categorized by levels of homology, topology, and distance between breakpoints. Error bars represent one standard deviation calculated by the bootstrap method, and stars indicate significant differences from the first subgroup of each category (p<0.0001).

The rate of rearrangements between any two loci is highly correlated with the rate at which each locus connects to other genomic loci (after controlling for rearrangement span; p < 10^-10^ for all spans; **Supp. Fig. 6**). For example, frequent rearrangements at chromosome 12 reflect juxtapositions to loci across the genome (**Fig. 1b**).

Rearrangements with high junction microhomology are substantially enriched in the cancer genome. Overall, for rearrangements with junction microhomology between 3 and 10 bp, each additional base of microhomology is associated with an approximately two-fold reduction in the number of rearrangements observed (**Fig. 2c**), a slower drop rate than the approximately four-fold reduction expected by chance^12^. We suspect that many of these rearrangements are due to microhomology-mediated end-joining (MMEJ); the effect of additional base-pairing on MMEJ has been confirmed in budding yeast^13^. For rearrangements with longer microhomology (>11 bases), we find each additional base of microhomology is associated with only a 7% decrease in the number of rearrangements observed. This transition may reflect a shift to single-strand annealing (SSA). In yeast, the transition from MMEJ to SSA appears to occur at 12-13 bp^13^.

Although rearrangements with high microhomology are enriched, as much as 70% of somatic rearrangements in our cohort are likely to be non-microhomology mediated. We determined this number by assuming that repair mechanisms that are not microhomology-mediated generate a random distribution of observed lengths of microhomology^12^ (**Ext. Fig. 2**).

We next compared microhomology levels across topologies, using the topological classification from Li *et al* (https://doi.org/10.1101/181339). We distinguished single, “isolated” rearrangements that were distant from other rearrangements from clusters of multiple “linked” rearrangements that represented more complex events. Among rearrangements with low microhomology (≤11 bp), isolated rearrangements tend to be short, whereas linked events tend to be long (**Fig. 2d**). Both isolated and linked rearrangements with high microhomology (>11 bp) are enriched with short events. The relationship between complex rearrangement topology and non-homologous or microhomology-mediated repair processes has been proposed in the context of chromothripsis^2^ and DNA replication mechanisms^14^ and may explain some of this observed bias.

We also observed significant enrichments for rearrangements connecting two genes and rearrangements connecting two repetitive elements, particularly for rearrangements connecting either two different long-terminal repeat (LTR) transposons or short-interspersed nuclear elements (SINE; **Fig. 2e**; **Ext. Fig. 3; Supp. Fig. 6**). Repetitive elements are sources of instability in the genome, and neighboring *Alu* elements have been reported to be sites of frequent recombination in the human genome^15,16^. Rearrangements joining repetitive elements from the same family exhibited significantly higher junction microhomology than rearrangements with only a single breakpoint within a repetitive element (**Fig. 2f**), suggesting that the former are enriched due to microhomology-mediated repair.

We used this information to develop two mathematically simple background models for the 2D analysis (**Ext. Fig. 4**; see **Methods**) that explicitly account for both the span distribution and the frequency with which each locus suffers rearrangements. The first model hypothesizes that the background probability that loci *i* and *j* will be juxtaposed is *p*^a^_ij_ = *q*_i_*s*_ij_ + *q*_j_*s*_ji_, where *q*_i_ is the marginal probability of a rearrangement initiated in locus *i* and *s*_*ij*_ is the conditional probability that a break at *i* will connect to site *j*. This “break-invasion” model is reminiscent of mechanisms like non-allelic homologous recombination (NAHR), which involve a break in one locus followed by invasion into another^17^. The second model hypothesizes the background probability *p*_ij_^b^ = *r*_i_*r*_i_*l*_ij_, where *r*_*i*_, *r*_*j*_ are the breakpoint densities and *l*_ij_ is a length factor connecting *i* and *j*. This “double-break join” model is reminiscent of non-homologous end joining (NHEJ) or MMEJ^18^, which involve separate breaks in two loci with an erroneous join.

The extent to which different classes of rearrangements fit either model therefore indicates the physical process that generated those rearrangements. We tested rearrangements stratified by level of homology, topology, and span (**Fig. 2g**). Rearrangements with no junction homology (<2 bp) show preference for the double-break join model, but rearrangements with increasing homology show increasing likelihood to be represented by the break-invasion model. Rearrangements whose junctions included an insertion longer than 10 bp (independent of junction homology), a characteristic often attributed to microhomology-mediated break-induced replication (MMBIR)^19^, were 10% more likely to fit the break-invasion model (p<10^-4^). Simple rearrangements tended to fit the break-invasion model whereas complex events tended to fit the double-break join model (p<10^-4^). Rearrangements shorter than 1 Mbp tended to fit the break-invasion model, whereas longer and interchromosomal rearrangements tended to fit the double-break join model (p<10^-4^ in all cases).

### Significantly recurrent breakpoints reflect multiple selective processes

The 1D analysis identified 52 loci where breakpoints were observed at rates significantly above the predictions of the background model, covering 38 Mbp or 1.2% of the genome (**Supp. Table 2**). The median size of a locus was 501 Kbp (s.d. 794 Kbp). Among the 2,693 genomes, 1,524 (57%) contained at least one SRB. The most significantly altered loci were sites of oncogenic fusions and regions surrounding recurrent somatic copy-number alterations (SCNAs), including a small number of fragile sites. From the top twenty recurrent loci, (**Fig. 3a**), five contained genes that are recurrently amplified (*TERT*, *ERBB2*, *VMP1/MIR-21*, *CCND1*, *MDM2*), four overlapped recurrent deletions of known tumor suppressors (*CDKN2A*, *PTEN*, *TP53*, *RB1*), seven contained genes involved in known oncogenic fusions (*TMPRSS2*, *ERG*, *BRAF*, *IGH*, *KIAA1549*, *BCL2*, *RUNX1*), and four involved genes at known fragile sites (*FHIT*, *WWOX*, *LSAMP*, *PTPRD*). Most SRBs were observed across several tissue types, but several of the known oncogenic fusions were identified in only a single tissue type.

**Figure 3:**
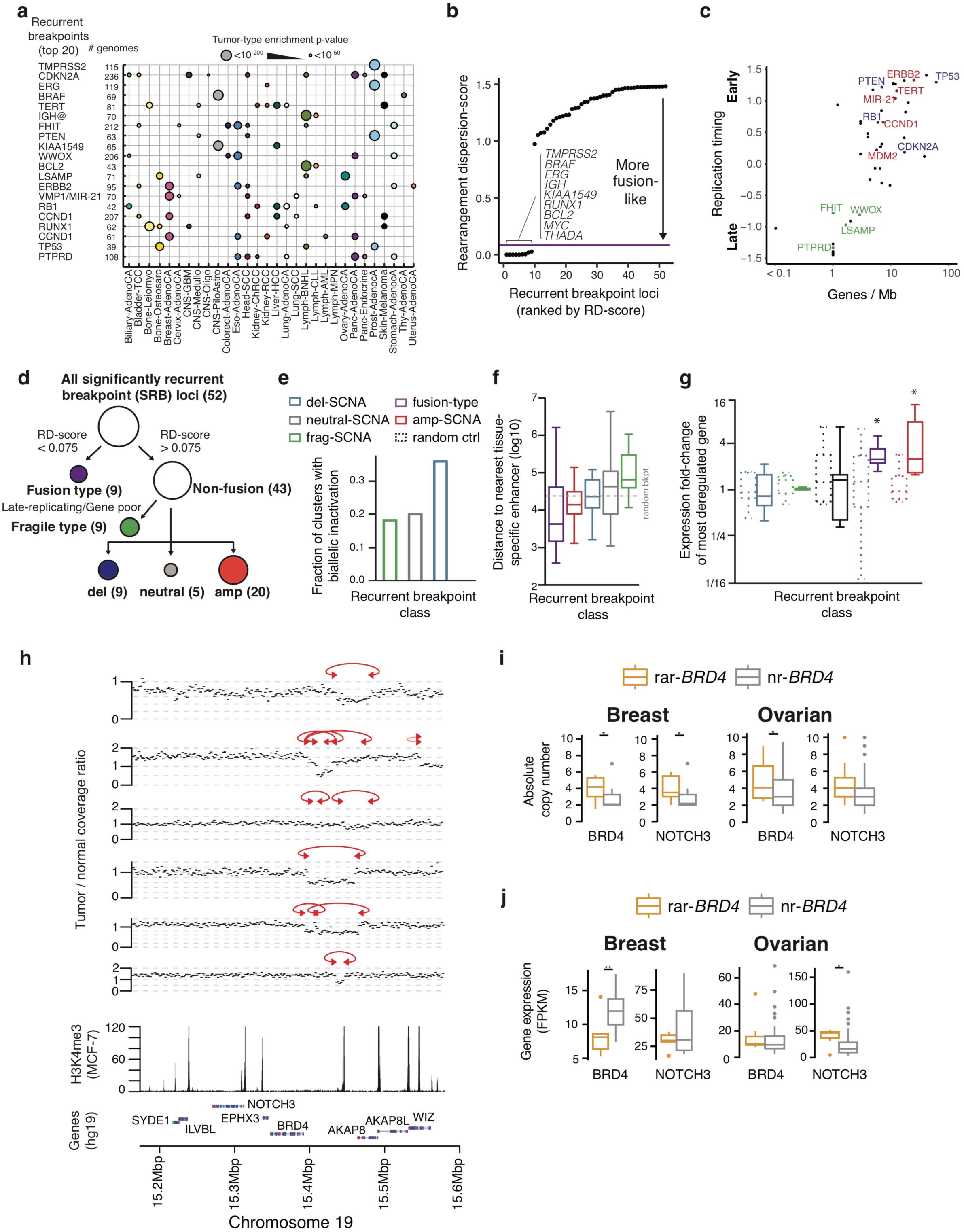
Significantly recurrent breakpoints (SRBs). a) The relative enrichment for events per histologic subtype (x-axis) for the top twenty most significantly rearranged loci (y-axis) is indicated by the size of the circles displayed. b) Ranking of RD-scores, representing the median absolute deviation of the distance between breakpoints relative to the median distance between breakpoints, for the 52 loci with SRBs. c) Gene densities (x-axis) and replication timing (y-axis) for recurrent breakpoint loci that were not classified as fusions. Known fragile sites (green) and driver SCNAs (blue: deletion; red: amplification) are annotated for the top 20 loci. d) Classification of recurrent breakpoint loci using RD-score, gene density, replication timing, and T/N coverage ratios. e) Fraction of recurrent breakpoint loci associated with biallelic inactivation of a known tumor suppressor gene. f) Distance in bp to the nearest tissue-specific enhancer (y-axis) for each breakpoint class. Dashed grey line represents randomly selected breakpoints. g) Expression fold-change (y-axis) for the gene with the most altered expression within 1 Mbp of the cluster centroid compared to samples without a breakpoint at the cluster locus. Random controls (in dashed boxes) represents randomly selected breakpoints. h) SRBs near *BRD4* in breast ductal and ovarian adenocarcinomas. i) Gene expression in FPKM (y-axis) for *BRD4* and *NOTCH3* in breast and ovarian tumors with (rar-*BRD4*) and without (nr-*BRD4*) *BRD4* rearrangements.

The rearrangements at the most significant loci exhibited two broad patterns (**Ext. Fig. 5**). In the first, the rearrangement breakpoints were clustered at one end, but the partner breakpoints were widely dispersed. These were largely associated with recurrent SCNAs. The second pattern involved rearrangements where both breakpoints were tightly clustered, resulting in oncogenic fusions.

With these patterns in mind, we developed a simple metric to quantify how tightly breakpoints cluster within each locus, to better group loci by their potential functional effects. For each locus, we calculated a “rearrangement dispersion-score” (RD-score) as the median absolute deviation of the distance between breakpoints, normalized by the median distance between breakpoints. The RD-scores of the 52 SRBs exhibited a bimodal pattern with a local minimum at 0.075. Nine loci had RD-scores below 0.075, with each locus containing genes involved in known oncogenic gene-gene fusions (e.g. *IGH*). We therefore classified these loci as “fusion-type” (**Fig. 3b**). The other 43 loci had RD-scores above this threshold and were associated with recurrent SCNAs and novel loci.

Among SCNAs, a major difficulty has been distinguishing recurrent alterations that are primarily driven by genome fragility from those resulting from positive selection^10,20^. We sought to improve this distinction by taking into account the rearrangements that generate these SCNAs. Fragile sites are thought to be generated from replication errors, and are associated with late-replicating^21^ and low gene-density regions of the genome^10^. We therefore scored the non-fusion loci by their gene density and replication timing, and found that the loci clustered into two distinct groups (**Fig. 3c**). The late-replicating, low gene-density group comprised nine loci, including each of the four known fragile sites among the top twenty loci. We therefore term these “fragile type” events. The remaining 34 non-fusion loci included known driver SCNAs and several loci not currently known to be altered by recurrent SCNAs. We therefore segregated these remaining 34 loci into those with significantly elevated copy-number (‘amplifications’, n=20), those with significantly decreased copy-number (‘deletions’, n=9), and copy-neutral events (n=5) (**Fig. 3d**; **Supp. Table 2**).

Although both fragile type and deletion events were associated with copy-loss, they appear to have different functional consequences. In particular, the rearrangements in more than one third of the deletion clusters were associated with biallelic inactivation of known tumor suppressor genes, whereas only 20% of rearrangements in fragile type clusters were (p<0.001; **Fig. 3e**).

The five rearrangement classes (fusions, fragile, deletions, neutral, amplifications) also had varied impact on the expression of genes in the immediate vicinity of their breakpoints. For each class, we compared the distance from the SRB to the gene with the most altered expression in the rearranged tumors, within a 1 Mbp window. Fusion-type clusters had the shortest distance to the nearest tissue-specific enhancer (**Fig. 3f**), and the change in expression of the most expression-altered gene was significantly greater than for deletions, neutral loci, fragile sites, and random nonsignificant breakpoints (p<0.05 in all cases; **Fig. 3g**). Of the three cluster types, fragile clusters displayed the weakest correlations with expression of neighboring genes, in agreement with the hypothesis that these are often passenger events that are not under selection^20^.

Many of the SRBs indicate novel and potentially functionally relevant events. For example, we observed recurrent deletions on chromosome 19 just upstream of *BRD4* and *NOTCH3*, which were significantly enriched for rearrangements in ovarian (10 tumors, p < 10^-8^) and breast (6 tumors, p < 0.006) adenocarcinomas (**Fig. 3h**). *BRD4* is a chromatin regulator and a candidate target of BET-bromodomain inhibitor therapy in several cancer types^22,23^, including ovarian and triple-negative breast cancer^24,25^. *NOTCH3* is located 36 Kbp away from *BRD4* and its activation may play a role in ovarian cancer^26^-^28^. The rearrangements in this locus tended to create tightly clustered <50 Kbp deletions near the *BRD4* promoter in both ovarian and breast adenocarcinomas. These rearrangements also tend to occur in cancers with amplifications of *BRD4* and *NOTCH3* (**Fig. 3i**), but are highly focal and do not contribute to those amplifications. Ovarian cancers, but not breast cancers, with these rearrangements had slightly increased expression of *NOTCH3*, consistent with its amplified state (**Fig. 3j**). However, *BRD4* expression was significantly decreased in the breast tumors and exhibited no change in the ovarian tumors, despite having increased copy-number in both tumor types. These findings, coupled with the expected result of disrupting the BRD4 promoter, raise the possibility that this cluster of rearrangements reduces *BRD4* expression in cancers where it would otherwise have been overexpressed. Overexpression of BRD4 has previously been found to suppress cell growth^29^. Deletions of promoters to prevent gene overexpression in the context of amplification has to the best of our knowledge not previously been reported in cancer.

### Significantly recurrent juxtapositions exhibit tissue-specific effects on expression

The 2D analysis (see **Methods**) identified 90 SRJs (juxtaposition clusters that were significantly enriched above expected rates; **Supp. Table 4**). Among the 30 most significant SRJs (**Fig. 4a**), 12 correspond to known oncogenic SRJs as curated by the COSMIC database (http://cancer.sanger.ac.uk/cosmic). An additional two clusters have been recently described to be oncogenic: a recurrent t(2;7) translocation between *THADA* and *IGF2BP3* in thyroid adenocarcinoma^30^ (in all cases we list the 5’ end of the SRJ first) and a recurrent t(22;23) translocation between *BEND2* and *EWSR1* in pancreatic endocrine tumors^31^. The sixteen remaining clusters include five with a known driver gene in the COSMIC cancer gene census (*MDM2*, *EGFR*, *TERT*, *ROS1*, *ERCC5*). Eight loci (*TMPRSS2*, *ERG*, *ROBO2*, *BRAF*, *TERT*, *BASP1*, *NEDD4L*, and *IGH*) were involved in more than one SRJ, a significantly higher number than would be expected by randomly choosing from all loci genome-wide (*p* < 10^-4^, permutation test), indicating different SRJs often share common molecular targets.

**Figure 4:**
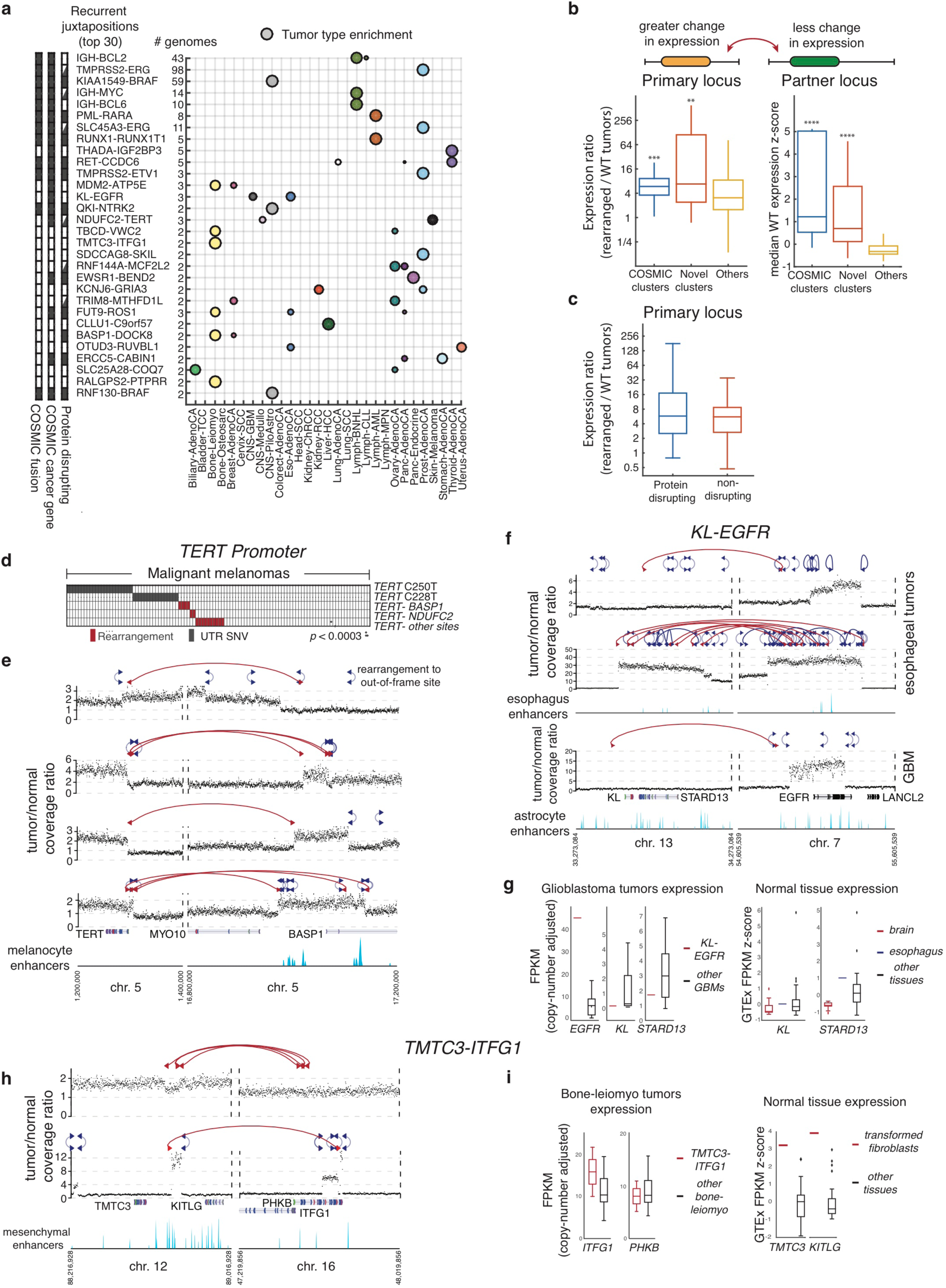
Significantly recurrent fusions. a) The thirty most significantly recurrently fused loci (y-axis), displayed by the relative fraction of events per histological subtype (x-axis). Fusion clusters are annotated by whether they are in the COSMIC list of oncogenic fusions (left bar, black box if on the list), whether at least one of their breakpoint loci overlaps a gene on the COSMIC cancer gene list (center), and whether all (solid black) or some (triangle) of the rearrangements within a cluster fall within introns or exons. b) Expression correlates for fusions in clusters in the COSMIC list (blue), other clusters (red), or not in any cluster (yellow). Displayed on the left is fold expression enrichment of the most highly overexpressed gene in the primary locus in cancer samples with these fusions relative to cancers of the same histology without the fusion. The primary locus is defined as the fusion breakpoint within 100 kb of the gene that is most highly overexpressed in samples with the fusion. Displayed on the right is the median expression level, in cancer samples of the same tissue type but without the fusion, of the gene closest to the partner locus breakpoint. Expression levels for the partner locus represent z-scores calculated across all cancers without the fusion. c) Fold expression enrichment of the most highly overexpressed gene in the primary locus, for fusions that disrupt protein-coding sequences (blue) and fusions that do not (red). d) Comut plot indicating *TERT* promoter mutations and rearrangements across all melanomas in the dataset. Promoter mutations and rearrangements were mutually exclusive. e) Rearrangements between *TERT* promoter and *BASP1* and *MYO10* locus result in focal amplification of *TERT* and relocation of enhancers to its promoter region. f) Recurrent translocation between *EGFR* in chromosome 7 and the *KL*/*STARD13* locus on chromosome 13. In all three samples the rearrangement contributed to the amplification of *EGFR*. g) EGFR expression in GBM tumor tissue after adjusting for copy-number (left) and KL and STARD13 expression in normal tissues (right). The single sample with the rearrangement and expression data showed high *EGFR* even after copy-number adjustment. h) Recurrent translocation in two leiomyosarcomas between a locus bordering *TMTC3* and *KITLG* on chromosome 12 and a locus bordering *ITFG1* and *PHKB* on chromosome 16. i) Expression of *TMTC3* and *KITLG* in (left) sarcomas with and without the rearrangement, and (right) fibroblasts compared to other tissue types.

Strikingly, nine of the ten most significant clusters comprise rearrangements from only a single cancer type. This restriction to individual cancer types seems to be specific to SRJs and contrasts with the other two major modes of somatic genetic alteration: copy-number alteration and single nucleotide variation, where the 10 most significant SCNAs and SNVs were each observed in an average of 11.9 and 6.7 cancer types, respectively (**Supp. Table 5**).

The finding that SRJs tend to be cancer-type restricted indicates that they are uniquely shaped by the epigenetic state of the cells in which they are observed. The epigenetic features that lead to this tissue specificity could favor mechanisms that generate specific rearrangements (e.g. due to varying three-dimensional organization of DNA among tissues) and/or tissue-specific selection pressures (e.g. differences in the transcriptional effects of the rearrangement or selective advantage for lineage-specific oncogenes)^32,33^. We hypothesized that the SRJs underwent positive selection in large part due to tissue-specific effects on expression of proto-oncogenes^4,34^.

We found three pieces of evidence that, as a rule, SRJs are enriched for their tissue-specific effects on expression of one of the rearrangement partners. First, rearrangements involved in SRJs lead to significant overexpression of one rearrangement partner relative to randomly selected rearrangements (**Fig. 4b** left panel, *p*<10^-4^). Second, the rearrangement partner that is not overexpressed in the rearranged samples tends to have high expression levels in that tissue relative to other tissue types, suggesting that the rearrangement brings tissue-specific regulatory elements associated with this gene to its partner (**Fig. 4b** right panel, *p*<2e-9). Third, the distance to the nearest tissue-specific enhancers is smaller for SRJs than for rearrangements overall (**Ext. Fig. 6**, *p*<1e-6).

In many cases, the selective pressures favoring SRJs also involve generation of novel protein coding sequences (including truncated and chimeric proteins). To some extent, this is expected: 33% of the mappable genome is covered by introns and exons, so a randomly placed rearrangement has a 56% chance of having at least one breakpoint fall within an intron or exon, and indeed 56% of all rearrangements do so. However, the rate is higher among SRJs, for which 68% have at least one breakpoint within an intron or an exon (*p* < 10^-7^).

However, the generation of novel protein coding sequences is not a general feature of SRJs. Only eleven of the 30 most significant SRJs generate novel protein coding sequences in all affected samples. An additional six exhibit a mix of protein-disruptive and nondisruptive rearrangements (**Fig. 4a**), but the protein-disruptive rearrangements in these six cases always occur within the first two introns of the disrupted gene, which can leave most of the affected protein intact^35^. Moreover, SRJs that generate novel proteins exhibit similar changes in expression to those that do not generate novel proteins (p=0.4; **Fig. 4c**), suggesting that altering gene expression is a function of both classes of SRJs.

The effects of SRJs are exemplified by the most significant cluster without a known COSMIC fusion: t(2;7) translocations between *THADA* and *IGF2BP3* in five thyroid cancers. In all five cases, the rearrangements connected *THADA*, truncated between introns 27 and 32, to a region just upstream of *IGF2BP3* on chromosome 7, always in the sense direction (**Ext. Fig. 7a**). Although not described in COSMIC, *THADA-IGF2BP3* fusions were recently shown^30^ to lead to *IGF2BP3* overexpression, promoting transformation. In our analysis, *IGF2BP3* was the most highly overexpressed gene in samples with *THADA-IGF2BP3* fusions, possibly because *THADA* has the highest tissue-specific expression in normal thyroid tissue (**Ext. Fig. 7b**). *THADA-IGF2BP3* juxtapositions were also mutually exclusive with rearrangements involving *RET* and other mutational driver events in thyroid cancers (e.g. *BRAF;* **Ext. Fig. 7c**), and were anticorrelated with *RET* expression (**Ext. Fig. 7d**).

The finding that juxtapositions tend to be tissue-specific does not imply that the oncogenes they generate are tissue-specific. For example, we found two novel SRJs involving *TERT. TERT* is known to undergo recurrent amplifications in 14 cancer types^36^, recurrent promoter mutations in over 50 cancer types^37^, and promoter rearrangements in 16 cancer types^38^. In our analysis, we identify significant juxtapositions between the *TERT* promoter region and *BASP1* in 4 melanomas (*p*<10^-7^) and between the *TERT* promoter region and *NDUFC2* in two melanomas and one medulloblastoma (*p*<10^-8^). Among melanomas, these rearrangements are mutually exclusive to the C225T and C250T *TERT* promoter mutations (p<10^-3^, **Fig. 4d**). Examination of the *TERT-BASP1* cluster indicates that in all four samples the rearrangement is part of a complex event resulting both in focal gains of *TERT* and what appears as a relocation of enhancers in and adjacent to *BASP1* to the *TERT* promoter region (**Fig. 4e**). Similarly, the *TERT-NDUFC2* rearrangements in the two melanoma samples result in both focal amplification of *TERT* and relocation of enhancers within a TAD containing *NDUFC2* and *ALG8* to just upstream of *TERT* (**Ext. Fig. 8c**).

Most SRJs were identified in only two or three samples, but even at this level of recurrence were highly significant. For example, we identified a translocation between *EGFR* on chromosome 7 and a locus adjacent to *KL* and *STARD13* on chromosome 13 in two esophageal adenocarcinomas and one glioblastoma (**Fig. 4f**). The likelihood of such narrow specific sites on two different chromosomes being connected in three different samples is less than 10^-8^. Additional features of these rearrangements support an oncogenic role. First, in all three samples these rearrangements appear to contribute to focal amplifications of *EGFR*. Second, *EGFR* was overexpressed beyond the expected level based upon its copy-number status in the glioblastoma sample (the only one with RNA-seq data; **Fig. 4g**), suggesting the rearrangement juxtaposed active regulatory elements to *EGFR*. Third, such regulatory elements seem to be active in esophageal tissue, where *STARD13* has somewhat higher expression than in most other tissues (**Fig. 4g**). Fourth, the glioblastoma sample with RNA-seq data exhibited low expression of *KL* and *STARD13* relative to other glioblastomas^39^. Both of these genes have been proposed to act as tumor suppressors^40^. A second example is a translocation connecting a region between *KITLG* and *TMTC3* on chromosome 12 with a locus just downstream of *ITFG1* on chromosome 16, in two leiomyosarcomas (**Fig. 4h**). Again, the likelihood of such specific regions on two different chromosomes being connected in even two samples is less than 10^-8^. Both of these samples show overexpression of *ITFG1*, a conserved transmembrane protein that may interact with the PP2A pathway and play a role in cell adhesion^41,42^ (**Fig 4i**). Both *TMTC3* and *KITLG* have high expression in normal fibroblast tissue (z-score > 3; **Fig. 4i**) and harbor an enhancer rich genomic region, suggesting that the overexpression of *ITFG1* may be due to the relocation of enhancers to its promoter region.

### TAD-disrupting rearrangements increase expression of neighboring genes more than TAD-preserving rearrangements and reveal novel oncogenic events

We further investigated the effects of rearrangements on expression by examining their interaction with TAD structure. TAD boundaries form functional barriers separating enhancer-promoter interactions^43,44^, and rearrangement-mediated TAD disruption can lead to activation of oncogenes^4^. For this reason, relative to TAD-preserving rearrangements, TAD-disrupting rearrangements have been proposed to have a larger impact on gene expression^45^. However, there has been no systematic investigation of such effects.

We therefore assessed the impact of rearrangements on gene expression, segregating rearrangements according to their spans and whether they disrupt nearby TADs on gene expression. We used CESAM, an analytical framework that we recently developed to integrate breakpoints with gene expression data and tissue-specific enhancer maps, to identify rearrangements associated with enhancer hijacking^4,34^. We found that TAD-disrupting rearrangements had a significant positive effect on gene expression compared with TAD-preserving rearrangements (p<0.001). We also found that this difference was most pronounced for rearrangements of 100-400 Kbp and became insignificant for >500 Kbp rearrangements (**Fig. 5a**).

**Figure 5:**
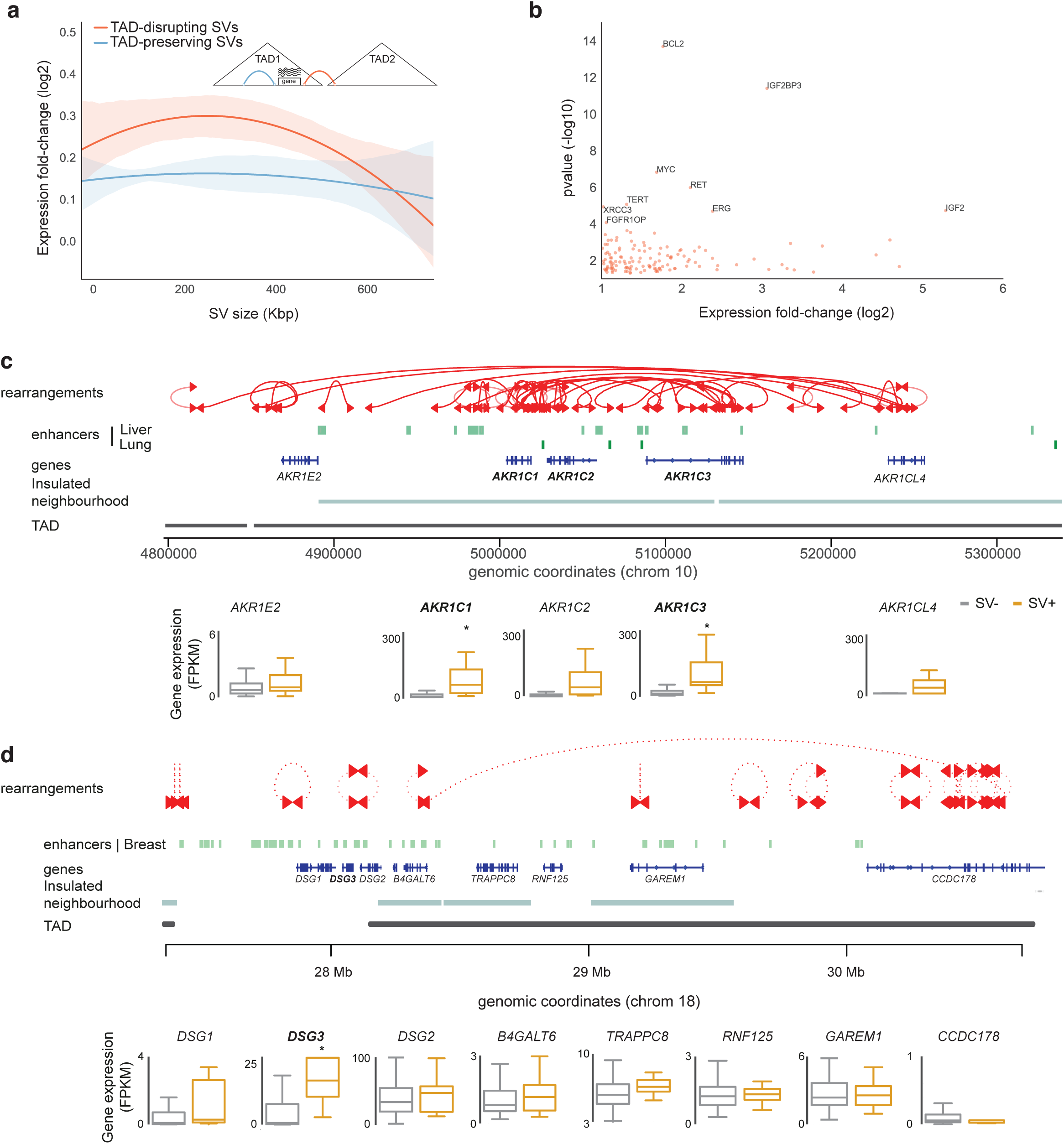
Impact and association of rearrangements on gene expression. a) Effects on expression (vertical axis) of TAD-disrupting and-preserving rearrangements, as a function of their span (horizontal axis). b) Volcano plot of significance (vertical axis) and associated gene-expression fold-change (horizontal axis) of CESAM hits for 1D, 2D and TAD-bound analyses. c) Schematic of rearrangements in the vicinity of *AKR1C* genes (top), locations of enhancers and TAD domains (middle), and expression of local genes (bottom) in samples with and without these rearrangements. d) Schematic of rearrangements in the vicinity of DSG genes on chromosome 18 (top), locations of enhancers and TAD domains (middle), and expression of local genes (bottom) in samples with and without these rearrangements.

We also performed a systematic search for specific rearrangements that disrupt TAD structures and are associated with strong (more than two-fold) changes in expression of nearby genes. We applied CESAM to TAD-bound breakpoint clusters for six sets of tumor samples, grouped by cell of origin (endoderm, mesoderm, ectoderm, neural crest, gastrointestinal and female urogenital organs) and evaluated all TADs for which at least three samples exhibited intra-TAD breakpoints and for which gene expression data were available.

Out of a total of 605 TAD-bound regions with sufficient data for CESAM analysis, we identified 190 TAD-bound regions whose disruption was associated with significant dysregulation of at least one gene (177 exhibiting upregulation and 13 downregulation; **Supp. Tables 3 and 6**). These included all 7 of the 54 SRBs and all 7 of the 90 SRJs for which we had sufficient number of samples with expression data to perform these analyses. Among these, 37 genes were classified as cis-activating events associated with enhancer juxtaposition. Many of the genes for which rearrangements were associated with upregulated gene expression were known oncogenes, including *BCL2*, *MYC*, *TERT*, and *IGF2BP3* (noted above).

Across the cell-of-origin groups, between 7% and 45% of affected TADs were associated with dysregulated gene expression, with tumors of neural crest and gastrointestinal origin displaying the highest proportions. The most highly upregulated CESAM hit associated with cis-regulatory rearrangement was a previously identified enhancer-hijacking event leading to *IGF2* upregulation^4^ in gastrointestinal tissues (mRNA expression 39-fold upregulated).

Several breakpoint clusters associated with robust expression alteration in *cis* could not be ascribed to previously described cancer gene loci. For example, we observed a breakpoint cluster at 10p15 (chr10:4.8 - 5.2 Mb), detected both in the 1D and TAD-bound CESAM analysis, which was associated with greater than two-fold upregulation of three *AKR1C* genes (*AKR1C1*, *AKR1C2*, and *AKR1C3*) within 11 Kbp of the breakpoint in seven lung squamous cell and two liver cancers (**Fig. 5c**). All breakpoints coincided with a cluster of lineage-specific enhancers at the locus, suggesting that the rearrangements may alter promoter-enhancer interactions at the locus to activate gene expression. Integration with chromatin conformation data revealed the *AKR1C*-family activating breakpoints to intersect with an insulated neighborhood^46^, which are three-dimensional topological structures that have been suggested to contain and ‘shield’ hard-wired enhancer-promoter interactions. All three *AKR1C* genes are within this insulated neighborhood and exhibited dysregulated expression in samples with these rearrangements. AKR1C proteins are aldo/keto reductases and involved in maintenance of steroid homeostasis. Ectopic expression of *AKR1C* genes can transform cell lines *in vitro* and germline mutations have been linked to increased susceptibility to lung cancer^47,48^.

We also identified an interesting pattern of rearrangements near *DSG3*, leading to upregulation in 13 breast cancer samples. Whereas the *DSG3* gene is situated at a TAD boundary, the rearrangements clustered inside this TAD, up to 2 Mbp away from the gene locus (**Fig. 5d** and **Supp. Table 3**). suggesting that these rearrangements perturb the TAD structure to activate *DSG3* gene expression. DSG3 is involved in cell-cell adhesion and has previously been implicated has a putative oncogene with a role in augmenting cell migration and invasion^49,50^, but the mechanism of upregulation has not been previously determined.

### Larger cohorts are required to detect SJRs that recur in up to 20% of cancers within individual tissue types

A central question in cancer genome discovery is how many samples need to be analyzed to detect recurrent driver events. We calculated the number of samples needed to obtain 90% power to detect SRJs as a function of the rate above background at which those SRJs recur and the distance between their breakpoints.

We first noted that more samples are required to detect short SRJs than long ones (**Fig. 6a**) due to the higher background rate (‘noise’) of the short events. For example, to detect a 100 Kbp SRJ that recurs in 0.5% of cancer samples (corresponding to 13 or more patients in our cohort) would require almost 3,000 samples–approximately the size of our pan-cancer sample set. (Reliable detection of such short SRJs will also require analytic improvements; see **Methods**.) Conversely, a 100 Mbp SRJ that recurs in 0.5% of cancers would require only slightly more than 1,000 samples.

**Figure 6:**
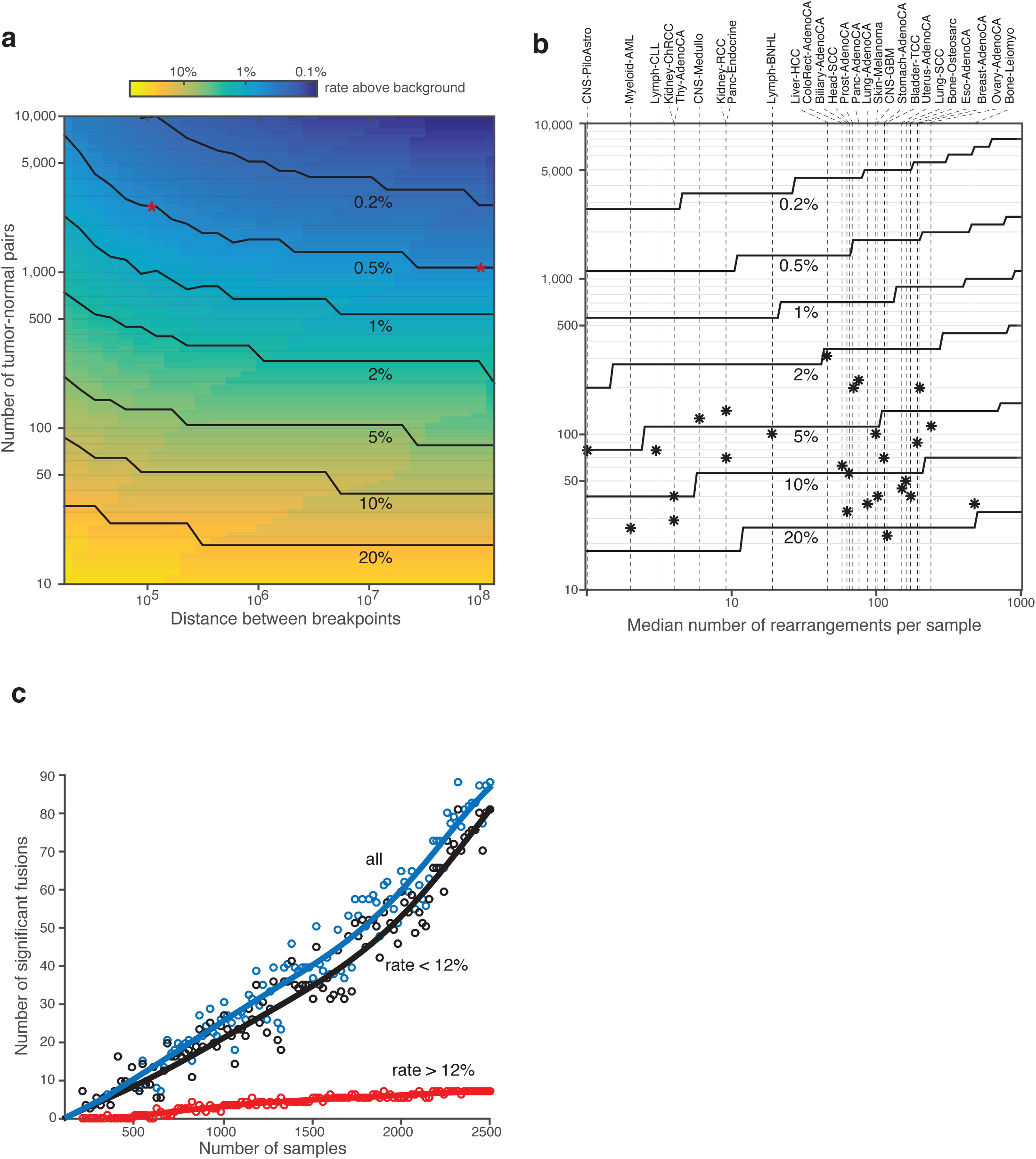
Power and saturation analysis of fusions. a) Number of tumor-normal pairs needed to detect fusions with 90% power as a function of the fusion’s span and the rate above background at which it recurs. The red asterisks indicate the numbers of samples required to detect 100 Kbp and 100 Mbp fusions that recur at 0.5% above their background rates. b) Number of samples (y-axis) required to detect 90% of recurrent fusions across 90% of pairs of loci, as a function of the median number of rearrangements per sample (x-axis) and the rate above background at which the fusion recurs (solid lines). The vertical dashed lines represent the median rearrangement rates of each cancer type, and the stars on these lines indicate the numbers of whole genomes analyzed for that cancer type. c) Number of significant fusions (y-axis) detected after down-sampling the data to various sample sizes (x-axis). Fusions that recur at high (>12%) and low (<12%) rates above background are indicated in black and red, respectively; their sum is indicated in blue.

We next integrated across rearrangement spans to determine how many samples are necessary to obtain 90% power to detect recurrent fusions across 90% of paired loci genome-wide (**Fig. 6b**, see **Methods**). We found that our pan-cancer analysis of 2,693 samples is limited to detecting rearrangements that recur in approximately 0.4% of all cancers.

However, SRJs tend to be tissue-specific, so we also calculated power using the number of samples available for each tissue type. We found that our current dataset, comprising 18-317 samples per tissue type (tissues with less than 15 samples were not considered), is powered to reliably detect 90% of driver fusions only for fusions that recur at a minimum of 2% (in liver cancers) to 21% (bladder cancers) above the background rate (**Fig. 6b**). For most cancer types, we are powered to detect events that recur in 5-20% of samples.

A down-sampling analysis of our data also indicated that we have not yet reached power to detect all recurrent driver fusions. When sufficient power is obtained (‘saturation’), reducing the number of samples modestly should not reduce the number of significant fusions clusters identified. However, when we down-sampled using random subsets with varying sample sizes, we found an additional novel SRJ for every additional 25 samples and no evidence of a plateau at sample numbers near the full dataset (**Fig. 6c**). SRJs that recur in greater than 12% of samples in a single tissue type did approach saturation, but SRJs that recur in fewer samples did not. For example, reducing the number of samples by 14% results in loss of detection of *QKI-NTRK2* fusions in pediatric gliomas–a potentially therapeutically relevant event^51^. The finding that low-recurrence SRJs are not approaching saturation suggests that adding more samples would uncover additional significant events.

## Discussion

The distribution of rearrangements in the cancer genome is shaped by both the mechanisms of their formation and the fitness advantages they confer on the cell. Our analysis revealed significant predictors of the distribution of rearrangement across the genome and identified known and novel rearrangements that recurred more often than expected given these predictions. Many of these recurrent rearrangements are likely driver events subject to positive selection, but it is possible some of them reflect mechanistic biases we did not account for. Indeed, nine of the SRBs likely reflect fragile sites in the genome.

Achieving a full understanding of the biological effects of recurrent rearrangements is complicated by the vast heterogeneity of their structure. We modeled rearrangements as both isolated breakpoints and as two-breakpoint juxtapositions. The vast majority of somatic rearrangements are in complex linked clusters, often involving several chromosomes (Li *et al*, cosubmission). Explicitly accounting for more complex topologies may improve our ability to detect driver rearrangements. Long-range connectivity information in the form of long read sequencing^52^, linked-reads^53^ and optical mapping techniques^54^ will be particularly useful for unravelling the structure of complex rearrangements. Such DNA sequencing should be accompanied by RNA sequencing to determine the expression consequences of these events.

Future efforts should be directed towards generating whole-genome sequencing data from many more cancers. The space of possible juxtapositions is the length of the genome squared, rather than simply the length of the genome, as is the case for other somatic genetic events. Large numbers of observed events are required to fill this space to understand the mechanistic biases influencing their distribution. Moreover, positively selected juxtapositions tend to be tissue-specific, and these are naturally more difficult to detect than alterations which span cancers. Our analysis indicates that we currently have sufficient power to detect fusions that recur in greater than 5-10% of cancer samples within each tissue type. However, we know that events that recur at lower rates can be biologically and clinically significant. For example, *ALK*-*EML4* fusions recur at a rate of 1-3% in lung adenocarcinomas (and only one sample in our cohort)^55,56^. At current sample numbers, we appear to be discovering a new novel fusion for every 25 cancer samples we sequence–a remarkable return on investment.

## Acknowledgements

Funds for this study were provided by the National Institutes of Health (T32 HG002295/HG/NHGRI, U54CA143798, U24CA210978, P01GM105473, R01CA188228, and R01CA215489), DFCI-Novartis Drug Discovery Program, Pediatric Low Grade Astrocytoma Foundation, the Cure Starts Now Foundation, Rigshospitalets Forskningsfond and the Danish Medical Research Council (DFF-4183-00233). We thank the members of the Technical Working Group of PCAWG for their assistance in generating the somatic mutation and rearrangement calls that contributed to these analyses.

## Extended Figure Captions

**Extended Figure 1:**
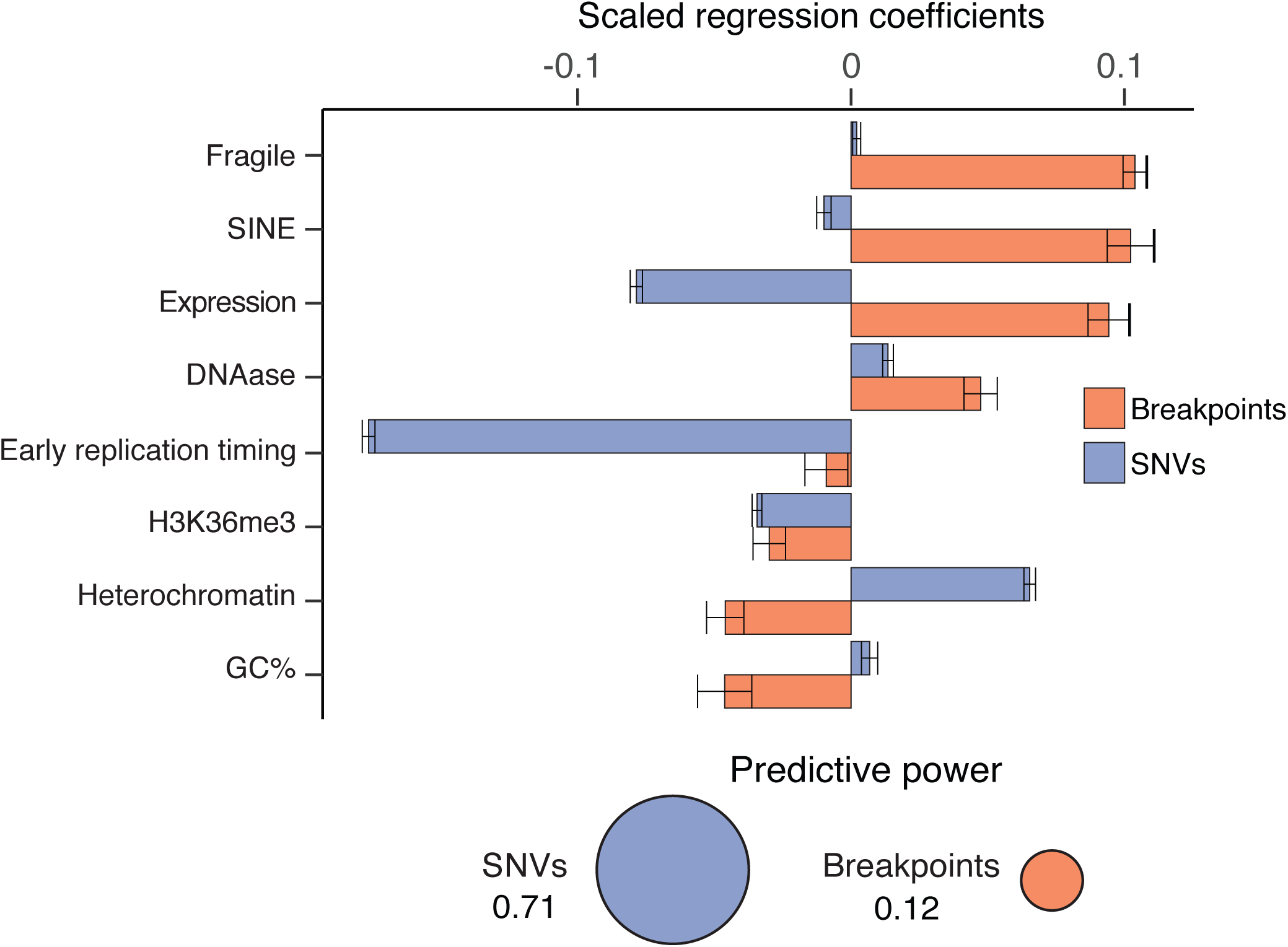
Genomic covariates predictive of rearrangement breakpoint frequency (orange) and SNVs (purple) using a Gamma-Poisson regression model. Regression coefficients less than zero predict a lower variant rate, and coefficients greater than zero predict a higher rate. The GP model explained 71% of the variability for SNVs and 12% for rearrangement breakpoints.

**Extended Figure 2:**
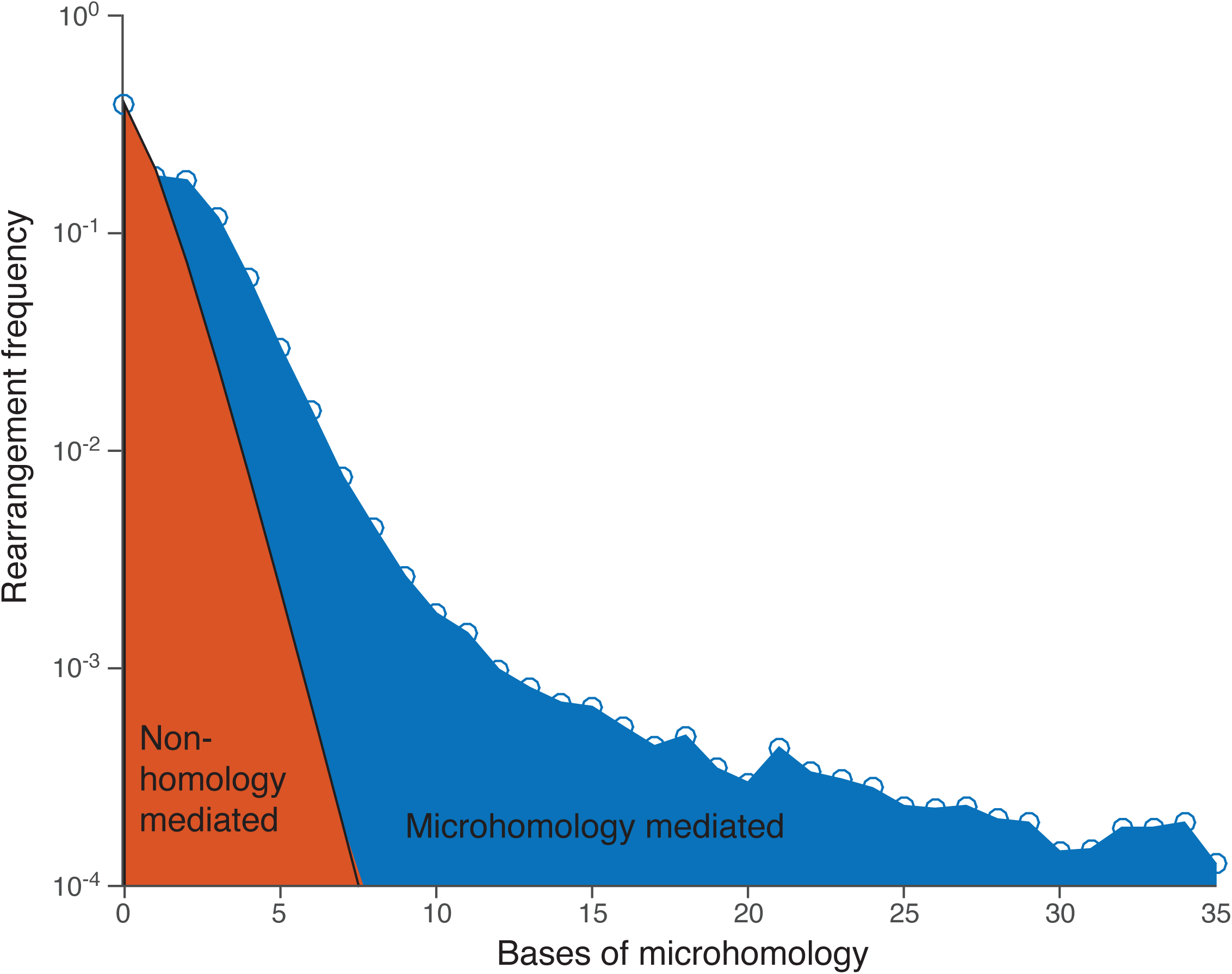
Genome-wide frequency of different levels of microhomology. Blue circles indicate observed data and black line indicates levels of microhomology expected by chance in non-microhomology mediated joining. The red region is attributed to NHEJ repair while the blue region corresponds to microhomology-mediated repair.

**Extended Figure 3:**
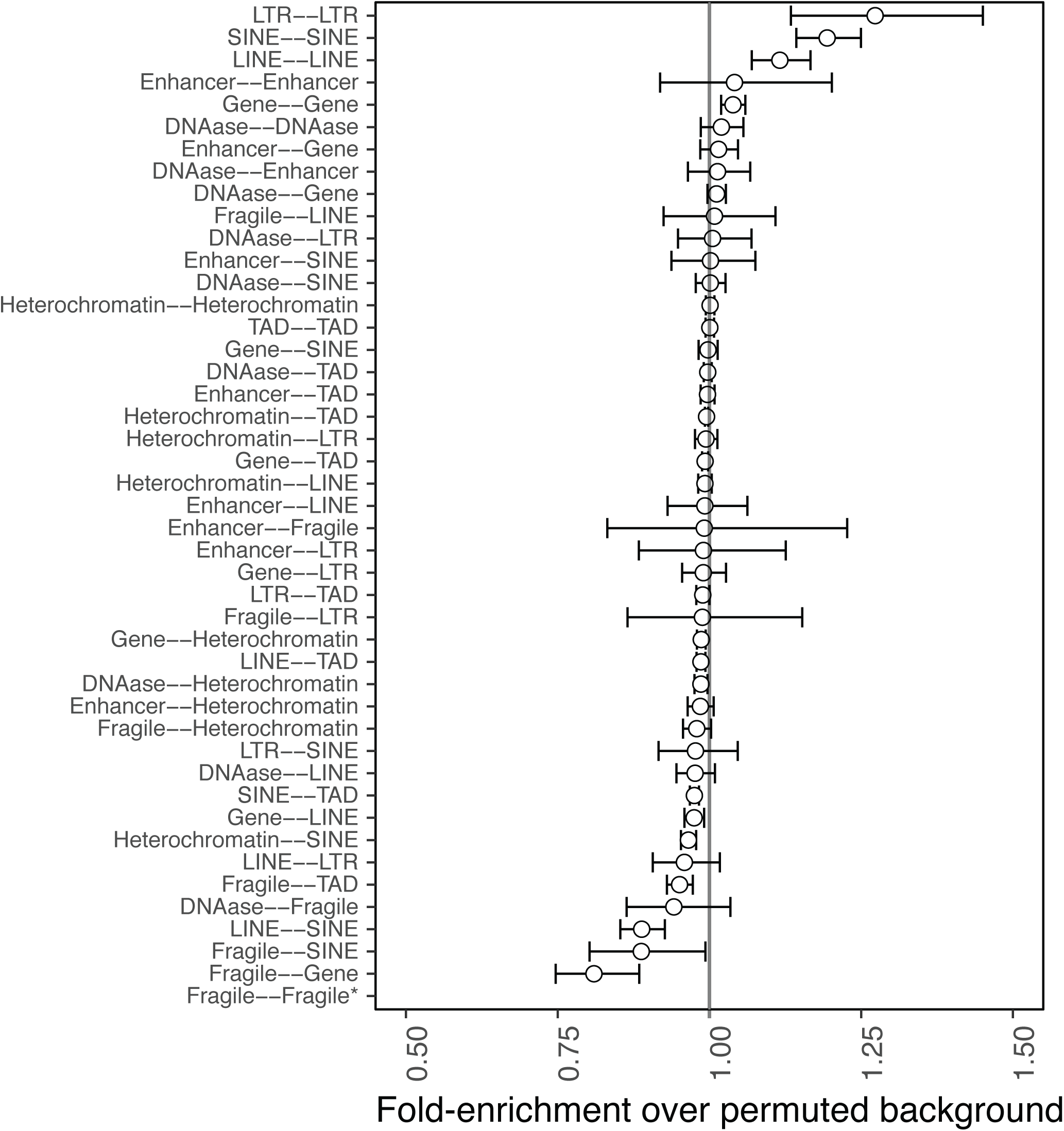
Enrichment or depletion for rearrangements fusing different genomic elements, compared with a permuted background. Enrichment scores near 1.0 (gray line) match a permuted background. Error bars represent three standard deviations. There were an insufficient number of fusions between different fragile sites to accurately assess an enrichment score.

**Extended Figure 4:**
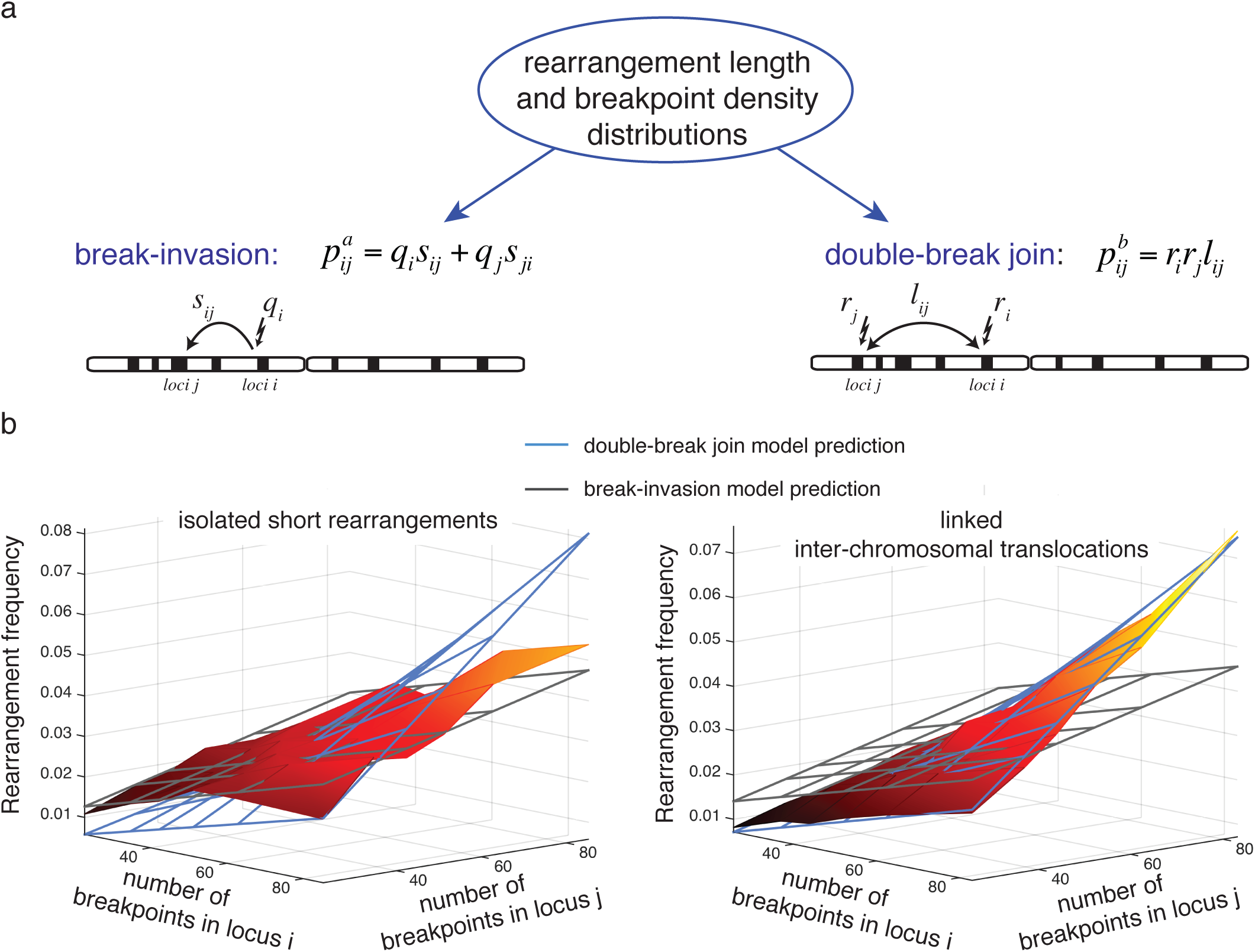
Two models for predicting the background rate of somatic fusions in the cancer genome. The break-invasion model (left) describes rearrangements that form at one locus (with probability q_i_) followed by invasion into another locus, with transition probability *s*_*ij*_. The double-break join model (right) describes rearrangements where both breakpoints occur independently (probabilities *r*_*i*_ and *r*_*j*_), and then fuse together with a factor *l*_*ij*_, which is a function of the distance between the breakpoints. b) The frequencies of rearrangements as a function of breakpoint densities, relative to model predictions. The observed frequencies of rearrangements are shown as a surface while the frequency predicted by the two models is indicated by solid blue and gray grids for the double-break join and break-invasion models, respectively. The left panel presents frequencies of isolated short rearrangements, and the right panel presents frequencies of linked inter-chromosomal rearrangements.

**Extended Figure 5:**
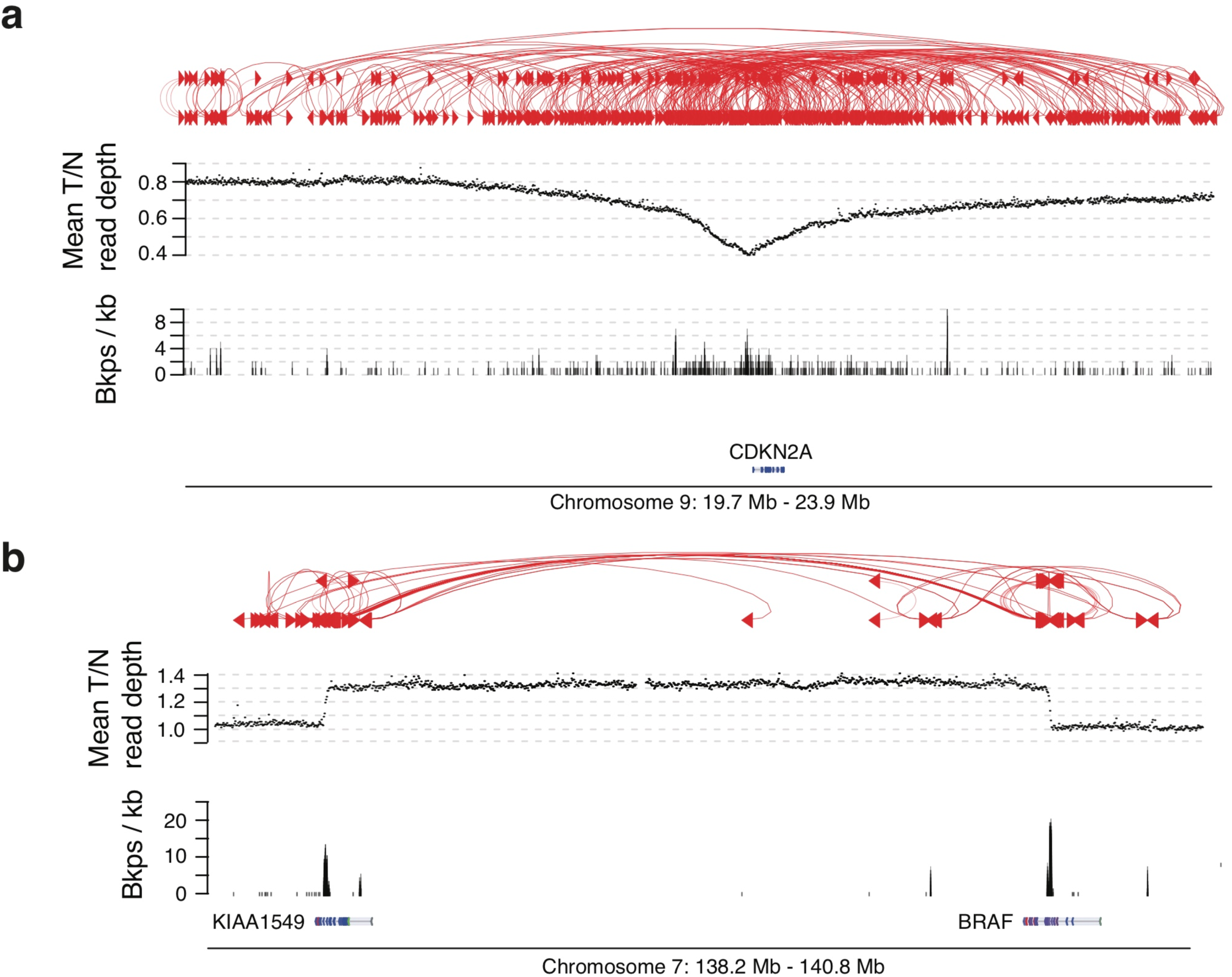
The two broad patterns of rearrangements observed at SRBs. a) Sites of recurrent SCNAs, such as *CDKN2A*, contain rearrangements whose partner loci are widely dispersed. b) Sites of recurrent juxtapositions, such as *BRAF-KIAA1549*, contain rearrangements whose partner loci are tightly clustered.

**Extended Figure 6:**
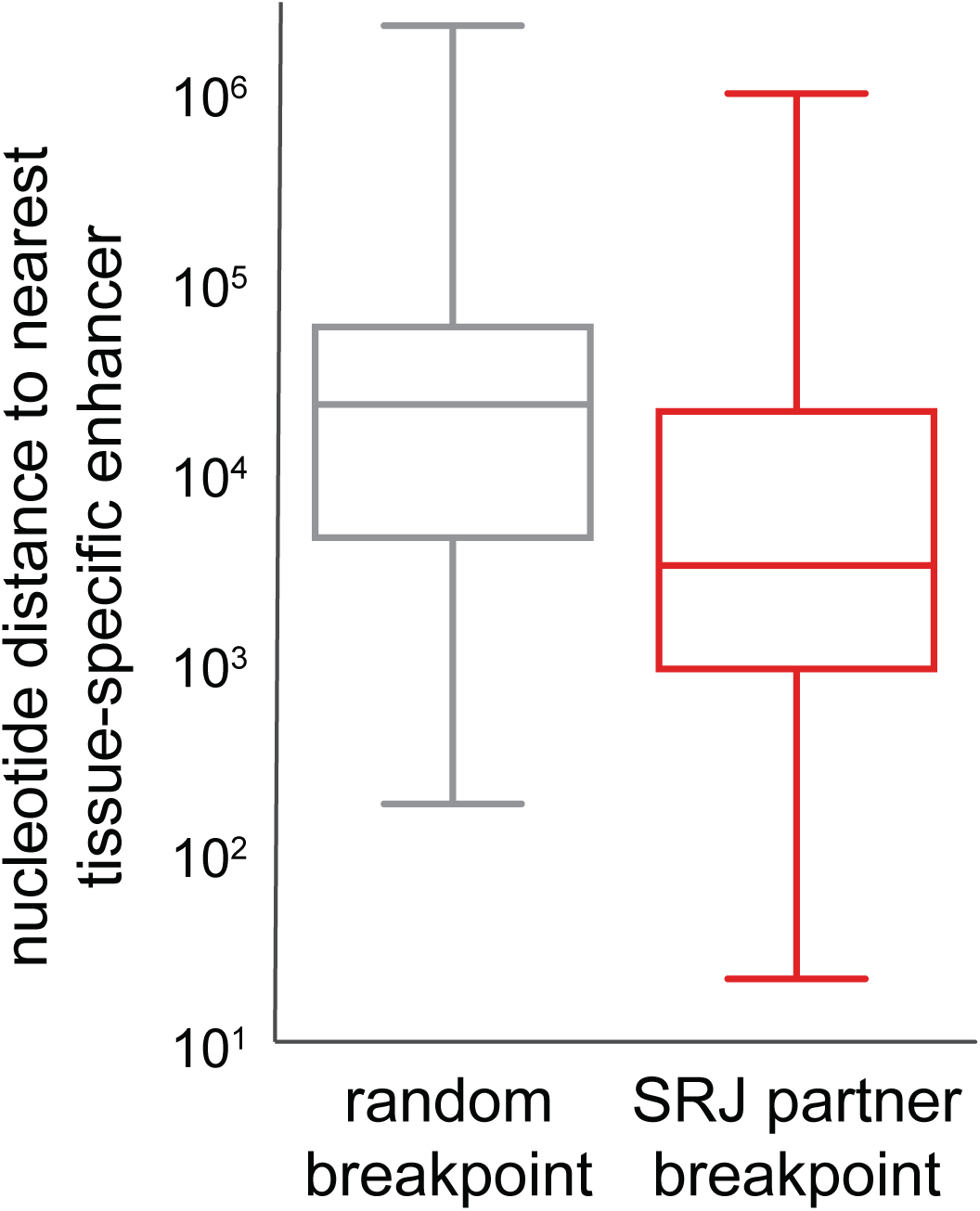
For SRJs, the distance from the partner site to the nearest enhancer is significantly smaller compared to randomly selected breakpoints.

**Extended Figure 7:**
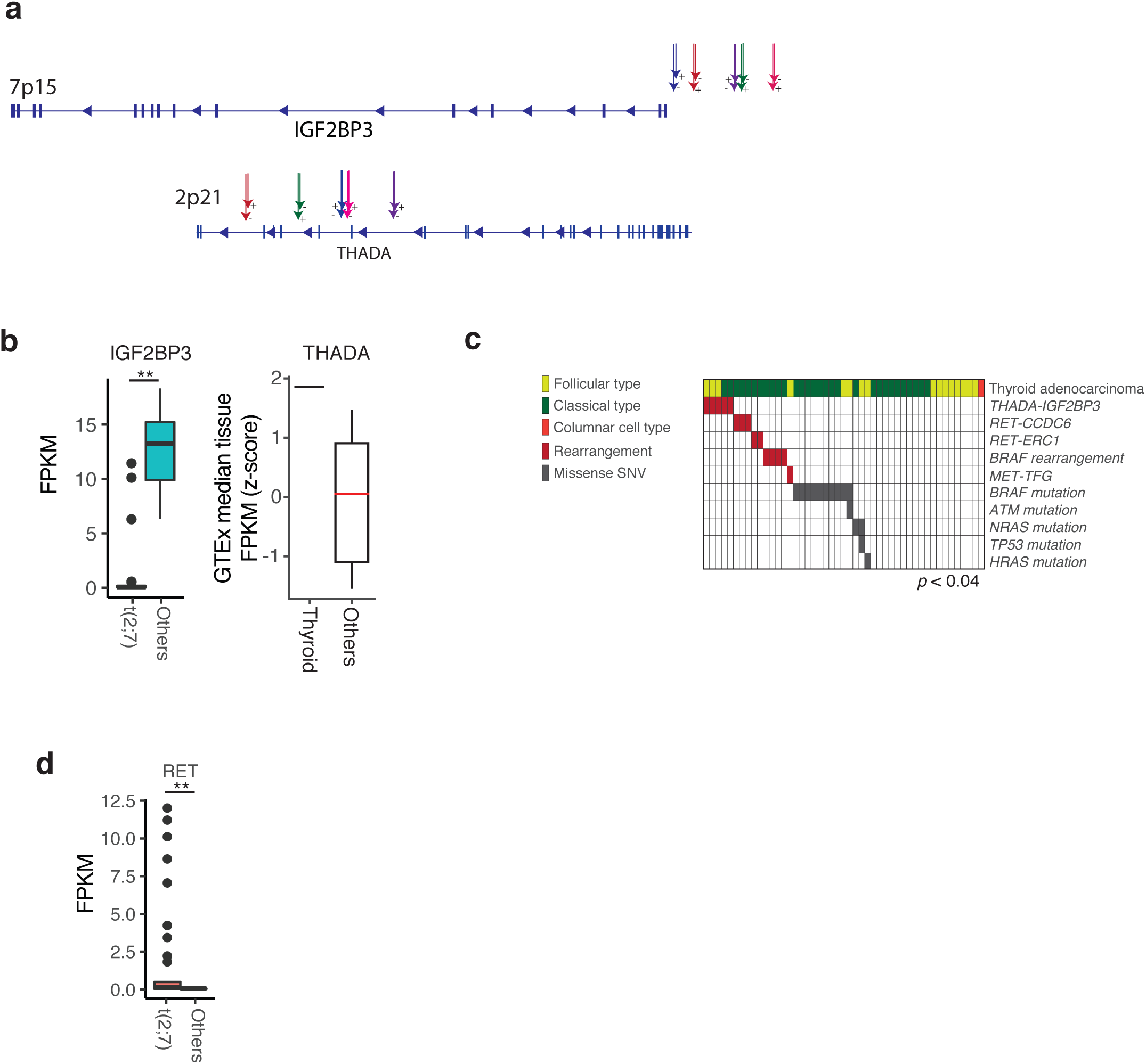
Recurrent t(2;7) fusions involving *THADA* and *IGF2BP3* in thyroid adenocarcinoma. a) Schematic of rearrangement b) Expression of *IGF2BP3* in thyroid samples with and without the rearrangement. Stars indicate a significant difference (p<0.01). c) T*HADA-IGF2BP3* are mutually exclusive with other known drivers of thyroid cancers, and d) are anticorrelated with RET expression.

**Extended Figure 8:**
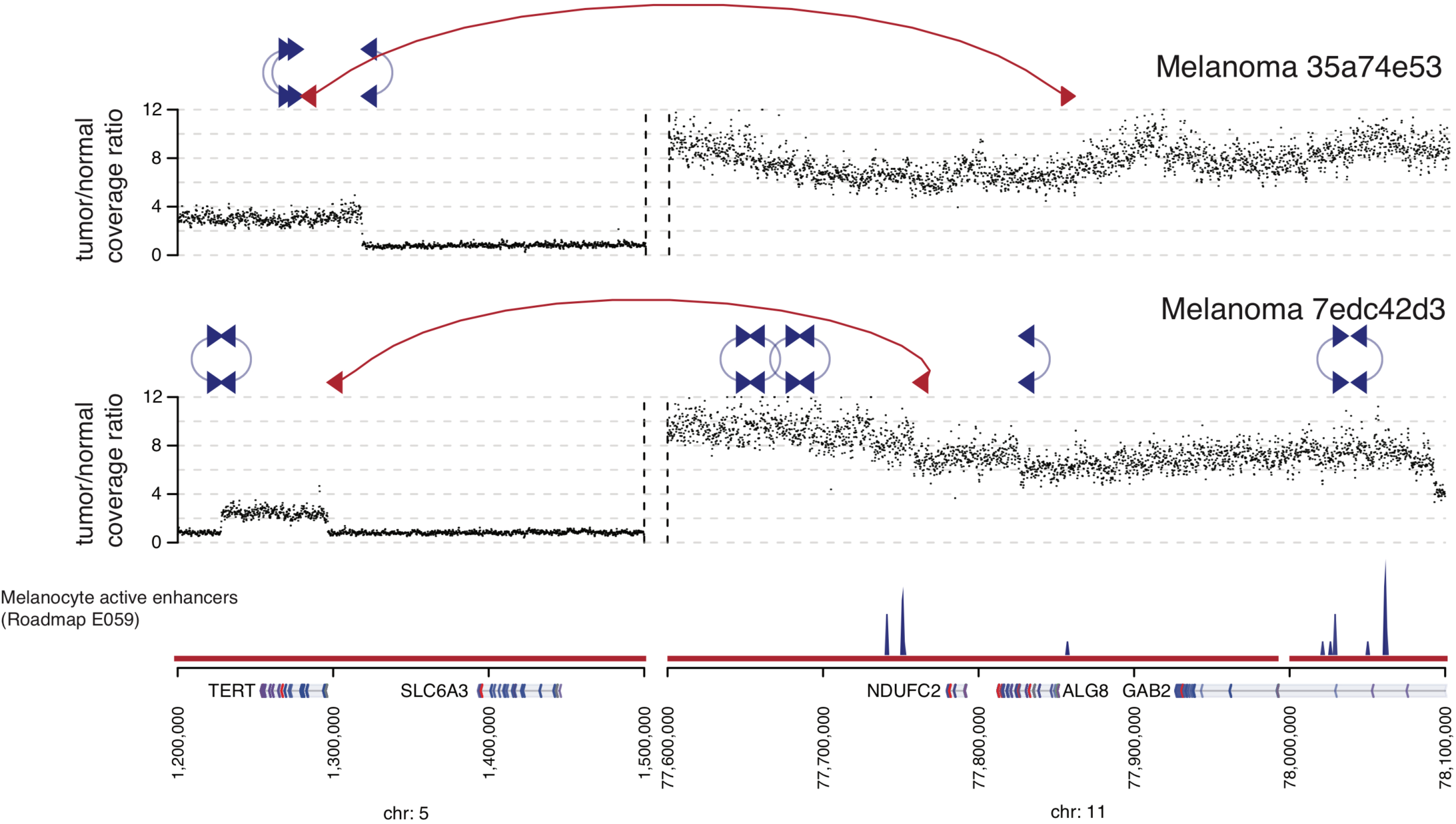
TERT-NDUFC2 fusion in two melanoma samples connecting *TERT* with an enhancer-rich region next to NDUFC2. Both samples also have focal amplifications of *TERT*.

## Supplementary Figure Captions

**Supplementary Figure 1:** The cohort of cancer genomes included in this study. a) A total of 2,693 cancer genomes were included, spanning 37 tumor types. b) Of the 2,693 donors with whole genome sequencing data, 1,241 had associated RNA-seq data (light blue).

**Supplementary Figure 2:** The merged rearrangement call set. a) Distribution of somatic rearrangements by tumor type. b) Comparison of the length of microhomology for somatic rearrangements, as called by SvABA (x-axis) and DELLY (y-axis). The area of the circle is proportional to the number of calls with a given microhomology. The two methods were largely consistent, with a zero-intercept linear model showing a slight trend towards higher microhomology calls from SvABA (perfect agreement: 1, observed: 0.928). c) Distribution of rearrangement call by support variant detection tool. d) Correlation between tumor cellularity and number of rearrangements (y-axis), by tumor-type (x-axis). There was no significant correlation between cellularity and number of rearrangements detected.

**Supplementary Figure 3**: Schematic of the germline CNV and class switch recombination filtering on the merged rearrangements calls. a) Germline CNVs identified by SvABA and present in 4 or more genomes from the PCAWG cohort were compared against the merged somatic rearrangements. Somatic rearrangements where each side overlapped the same germline CNV were reclassified as germline CNVs and excluded from further analysis. b) Example of filtering class switch recombination rearrangements at the IGH@ locus. Rearrangements that begin in IGH@ (purple, portion shown), IGK@ or IGL@ and connect to a different locus (e.g. BCL2, portion shown in green) are retained. Rearrangements with both breakpoints in the immunoglobulin loci are removed.

**Supplementary Figure 4**: Gamma-Poisson model for identifying signatures of mechanism and selection for rearrangement breakpoints. a) Histogram of breakpoint counts per 50 Kbp bin (with only one sample-breakpoint per bin). The distribution more closely follows a Gamma-Poisson (red) than a Poisson (green). b) Diagram of the full breakpoint model (red) and a naïve control with no covariates (green) and a randomized control with all covariates but shuffled around the genome (blue). The full model produces the highest log-likelihood (y-axis, bar chart), relative to the naïve and random models. c) Scaled regression coefficients (x-axis) for the three models using 11 covariates. Positive coefficients increase the predicted breakpoint count, negative coefficients lower the breakpoint count. The naïve model used only the mappability covariate. Error bars represent 95% confidence intervals.

**Supplementary Figure 5**: Quantile-quantile (QQ) plot of the probability that the breakpoint frequency within each 50 Kbp locus in the genome occurred at the observed rate or higher (y-axis) compared against a uniform probability distribution (x-axis). The genomic inflation factor (red) for this model was 1.09. An inflation factor much lower than or much greater than unity describes a poorly fit model.

**Supplementary Figure 6**: Regression plots of the relationship between the number of rearrangements connecting loci i and j and the breakpoint density at i (excluding rearrangements to j). Four distances between i and j are tested: a) span < 5x10^4^ b) 10^6^ < span < 5x10^4^ c) span > 10^6^ and d) inter-chromosomal translocations.

**Supplementary Figure 7:** Permutation model for identifying two-dimensional correlates of rearrangements. a) Schematic of the Swap method. A sparse matrix representing rearrangements [1] is permuted by swapping the *x* and *y* coordinates of two randomly selected points, perfectly preserving the breakpoint counts per locus. After a burn-in period, swaps that disrupt the rearrangement length distribution are rejected [4]. The full permutation scheme is applied *N* times to create *N* separate matrices [5]. Connections between two loci are represented as areas on the matrix, and rearrangements are tallied by their membership in different areas [5]. The histogram of rearrangement tallies (black) from the *N* permuted matrices are compared with the observed tally (red line) [6]. b) Euclidean distance between the permuted and empirical rearrangement span histograms (y-axis) as a function of the number of swaps (x-axis). After an initial period of 5 million swaps for inter-chromosomal translocations (no span dependence), intra-chromosomal swaps move the span distribution initially away from the observed empirical distribution. The swaps are then rejection sampled to accept only those that move the span distribution back towards the observed. c) The distribution of fusions in the PCAWG cohort (original) compared with the permuted distribution from Swap (permuted data). d) Swap results for true SINE elements (left), compared with randomized SINE elements (right). e) Same as (d), but restricted to only inter-chromosomal rearrangements.

**Supplementary Figure 8:** Binning scheme for the fusions recurrence analysis. Distributions of (left) bins sizes and (right) number of rearrangements per bin.

## ONLINE METHODS

### Annotation of potential functional effects of rearrangements

We annotated the potential functional effects of each rearrangement based on the locations and orientations of its breakpoints. Gene definitions for genome build hg19 were obtained from the UCSC Table Browser^57^. Breakpoints were annotated by whether they fell inside the body of a gene, and by all of the genes fully or partially contained within 100 Kbp of either side of the break. Intergenic breakpoints were annotated with the distance to the nearest gene.

Rearrangements were evaluated for whether they could produce a possible in-frame sense fusion transcript. The CCDS database^58^ from hg19 was obtained from the UCSC Table Browser. With the CCDS intervals, breakpoints contained within a gene were annotated by which intron or exon they overlapped with, and the coding frame (1,2, or 3) of the first exon opposite the direction of the breakpoints. Candidate fusions were called as in-frame and sense if 1) the relative orientations of the breakpoints and directionality of the gene resulted in a potential sense fusion and 2) the two breakpoints we in the same coding frame.

### Classification of rearrangements by topology

We classified the rearrangements into topological groups based on their orientations, spans, and whether they were significantly closer to neighboring breakpoints than expected by chance, potentially indicating a complex rearrangement. A detailed description for this classification scheme is described in our companion paper by Li *et al*. We combined the classifications by Li *et al* into five major groups (complex rearrangement clusters, simple deletions, simple inversions, tandem duplications and translocations) and examined their distribution across the genome.

### Assessing the significance of somatic rearrangement breakpoints

#### Modeling breakpoint counts with a Gamma-Poisson regression model

To model the background rate of somatic breakpoints, we first established a discrete coordinate system on which to evaluate genomic covariates and breakpoint counts. We binned the genome into 50 Kbp bins, with 1 Kbp of overlap between bins to reduce edge effects. Excluding gaps in the reference genome (hg19) and the sex chromosomes, this produced 50,561 loci. Complex events with many tightly clustered breakpoints could dominate the breakpoint count at a single bin and cause an over-estimation of the prevalence of breakpoints at those loci. To account for this, we only considered one breakpoint per donor per locus. After removing locus-donor duplicates, 336,496 breakpoints (55% of all breakpoints) were counted within our model. The number of breakpoints per bin ranged between 0 and 120, with a median of 5.0 and mean of 6.1. The vast majority of bins (99.0%) of bins contained 20 or fewer breakpoints, and 2.6% contained zero breakpoints.

The detected rate of breakpoints across the genome is also confounded by the mapping quality within a locus. Rearrangements in regions that are difficult to align to (e.g. alpha-satellite repeats) were rejected by our variant callers, leading to a relative depletion of events in regions with low mappability. To control for this effect, we use the concept of “eligible territory” from Imielinski et al^59^, and normalized the breakpoint counts within each locus by the number of bases eligible for breakpoint detection. To establish an eligible territory, we used the “universal mask” described in Li 2014^60^ and used in Imielinski et al^59^. Briefly, this mask filters regions of low mappability, low complexity and sites of unusually high numbers of aberrant SNV calls from the 1000 Genomes Project.

The distribution of breakpoint frequencies per bin was widely over-dispersed (D = 9.12), suggesting a Gamma-Poisson (GP) fit to the data (or equivalently, a negative binomial). Compared with a Poisson fit, the GP distribution produced a 1.1-fold higher log-likelihood (Supp Fig. 4a). We therefore elected to model the breakpoint frequencies using a GP regression model, where the log of the expected value of the breakpoint counts per bin could be modeled as a linear combination of genomic covariates within each bin.

We then applied a GP regression model for breakpoint count data, adapted from the model for SNVs and indels from Imielinski et al^59^, and specified as:

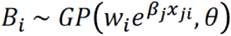

where w_i_ is the eligible territory of locus i, B_i_ is the breakpoint count at locus i, x_ji_ is the matrix describing the values of covariate j at locus i, and (theta) is a single scalar representing the shape parameter of the distribution. The regression coefficients (beta) were then found by maximum likelihood estimation using MASS∷glm.nb in R-3.3.2, which utilizes the NB2 parameterization of the GP function. The source code for the GP model is available at https://github.com/mskilab/fish.hook.

#### Genomic covariates that predict breakpoint frequencies

We hypothesized that local sequence features (e.g. density of repetitive elements), replication-timing, chromatin state, epigenetic modifications, and other genomic features, could be predictive of breakpoints rates within our GP model. We therefore fit our GP model using both “interval” covariates that indicate genomic regions (e.g. SINE elements), and “numeric” tracks that indicate values (e.g. GC content) associated with genomic regions. To enable direct comparisons between different covariates, each covariate was transformed to a z-score, centered at zero, using stats∷scale (R-3.3.2). The complete list of genomic covariates and their scaled covariate scores are for both the breakpoint and SNV models are listed in **Supp. Table 1**. We evaluated the effects of these covariates using three different GP models: the full model with all covariates, a naïve model using only the mappability covariate, and a random model using all covariates but with permuted annotations of genomic locations for each track. We then calculated the log-likelihoods of each model using stats∷logLik (R-3.3.2), and found that the full model achieved a significantly higher log-likelihood than the naïve or random models (**Supp. Fig 4b**). The random model scored very slightly higher than the naïve model, likely due the added degrees of freedom and possibility for over-fitting. However, relative to the full model, the difference was small, suggesting that the full model has a low degree of over-fitting. The naive and random models each explained 0.02% of the variance (Pearson R^2^) compared to 12% for the full model.

We note that several factors are limiting the variance explained of the full model. First, the training set of the regression model included loci that undergo selection which can not be excluded *a priori*, and therefore inherently limit the variance explained. This is in contrast to SNVs were synonymous mutations can be used to construct the background model. Second, in this work we study a set of covariates (Supp. Table 1), and it well may be that additional covariates and relationships (e.g. non-linear) would result in a better model. Third, heterogeneities in breakpoint rates, for example across different cancer types, may require independent treatment (addressed by Li *et al)*.

#### Assessing the significance of loci with high breakpoint rates

We used the full GP model to estimate the background rates for each locus and to calculate the probability that c_i_ or more events would be observed at locus *i*. The count data *c*_*i*_ is restricted to a non-negative integer, and the probabilities will be a slight over-estimate of the true value. To correct for this, we use the procedure employed in Imielinski et al^59^ to select a random probability from a uniform distribution between the probability of observing *c*_*i*_ breakpoints and the probability of observing *c*_*i*_ + 1 breakpoints. To correct for multiple hypothesis testing, we calculated the false discovery rate (FDR) using the Benjamini-Hochberg method^61^. At an FDR cutoff of 10%, we observed 206 separate 50 Kbp loci (0.4% of the tested loci) to be significantly enriched for breakpoints beyond the predicted rate.

We next attempted to determine which breakpoints at each locus under positive selection were themselves likely driver rearrangements. We noted that the breakpoint counts at many loci were dominated by rearrangements from a small subset of tumor types, suggesting that the rearrangements in these tumor types were drivers. Some rearrangements from other tumor types, however, would often also be seen at background rates expected for these tumor types.

We therefore calculated an enrichment p-value (binomial test) that tumor type *T* was enriched at that locus:

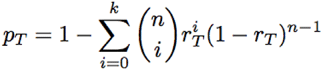

where *k* is the number of breakpoints from tumor type *T* intersecting the locus, *n* is the total number of breakpoints intersecting the locus, and *r*_*T*_ is the fraction of breakpoints from tumor type *T* within the entire PCAWG cohort. Using this enrichment score, we considered as driver rearrangements only rearrangements from the most enriched tumor-type and any tumor-type *x* with log(p_x_/p_top_) < 3. Analysis of the p-values distribution and quantile-quantile plot (**Supp. Fig. 6**) shows a uniform distribution without apparent biases of p-values.

The 206 significantly recurrent 50 Kbp loci tended to form clusters around a smaller number of distinct loci (e.g. around *CDKN2A*). We therefore merged these clusters together by joining significant loci and their intervening regions if they were separated by fewer than 200 Kbp. This reduced the 206 recurrent 50 Kbp bins in 52 significantly recurrent loci.

#### Comparison of recurrent breakpoint loci with significantly recurrent SCNAs and known fusions

We compared the significantly recurrent breakpoint loci with sites of significantly recurrent SCNAs obtained from GISTIC2^62^ analyses and available on https://www.synapse.org/#!Synapse:syn8341168 and the COSMIC cancer database curated list of gene fusions in cancer (http://cancer.sanger.ac.uk/cosmic/fusion). Recurrent breakpoint loci that overlapped a GISTIC peak region (deletion or amplification) from either the pan-cancer (all_cancers) analysis or any tumor-type specific analysis were considered as representing a recurrent SCNA. Recurrent breakpoint loci among the top twenty (ranked by q-value) were labeled according to their overlap with known cancer genes from the COSMIC cancer gene census (http://cancer.sanger.ac.uk/census). Recurrent fusions were considered supportive of a known fusion if the two loci involved in the recurrent fusion overlapped both genes from an entry in the COMIC fusion database.

### Classification of rearrangement patterns at sites of recurrent breakpoints

To predict the functional effects of the recurrent breakpoint loci, we scored each locus based on its pattern of rearrangements and genomic covariates. For each rearrangement containing a significantly recurrent breakpoint, we calculated the “RD-score”, which we defined as the median absolute deviation (MAD) of the breakpoint-breakpoint distance (or 10^9^ for inter-chromosomal rearrangements) divided by the median breakpoint-breakpoint distance. For inter-chromosomal rearrangements, we evaluated only rearrangements to the most frequent chromosome. Rearrangements at sites of known recurrent oncogenic fusions exhibited low RD-scores (e.g. *IGH-BCL2*, RD-score: 0.01), while breakpoints at known fragile and driver SCNA sites exhibited a high RD-score. Hartigans’ dip-test (in R v3.3 - diptest∷dip.test) supported a non-unimodal distribution (p = 0.02) with a discriminant of 0.075. The RD-score for all significant loci is listed in **Supp. Table 2**. Recurrent breakpoints with RD < 0.075 were classified as supporting fusion-type driver events.

For each recurrent breakpoint locus not classified as fragile-type or fusion-type, we evaluated whether the breakpoints tended to delete, amplify or leave unchanged (neutral) the copy-number state of the adjacent region. For each locus with width *W*_*i*_, we calculated the mean tumor-over-normal coverage ratio from the raw coverage tracks (binned to 2 Kbp). We then compared this with the mean tumor-over-normal coverage ratio for the region immediately to the left (width *W*_*i*_) and immediately to the right (width *W*_*i*_). Regions with significantly higher coverage profiles in the target window were classified as amplification-type, while regions with significantly lower coverage were classified as deletion-type.

### Identifying mechanistic factors influencing fusion partner selection

We developed Swap, a non-parametric model for identifying factors that influence the frequency with which two genomic locations will be fused (**Supp. Fig. 7a**). Swap controls for the frequency of breaks at a given locus and the span (l) dependence of rearrangements, which favors short events by a factor proportional to the inverse of the rearrangement span. Swap is available at https://github.com/walaj/ginseng.

Swap permutes the data by randomly choosing two rearrangements (x_1_, y_1_) and (x_2_, y_2_), with the requirement that they either both represent inter-chromosomal translocations, or both represent intra-chromosomal rearrangements. The x and y coordinates are then swapped to create the new rearrangements (x_1_, y_2_) and (x_2_, y_1_). This preserves the total number of breakpoints at a given locus (no new x or y coordinates are produced), while randomizing the fusions joining two breaks.

The random swaps tend to change the span distribution from the 1/l distribution towards a uniform distribution. To preserve the empirical span distribution, after an initial unconstrained set of permutations, Swap only accepts only intra-chromosomal swaps that move towards the empirical span distribution. To calculate the difference between the swapped and empirical span distributions, we log-transformed the span distributions and created a histogram with 20 equally spaced bins in log-space. After each swap we find the Euclidean distance *D* between the observed empirical histogram *H*_empirical_ and the histogram H_permuted_ from the permuted matrix (**Supp. Fig. 7b**). We continue randomly swapping points until the distance between the permuted and the empirical histogram is less than 5%. The final product is a permuted set of rearrangements that has the same distribution of spans and breakpoints as the original data (**Supp. Fig. 7c**).

To test the hypothesis that a rearrangement fuses loci A (e.g. all SINE elements) to loci B (e.g. all LINE elements), Swap compares the number of observed A-B fusions with the distribution of A-B connections in the collection of randomized matrices. The A-B enrichment or depletion factor is the ratio between the observed average number of permuted A-B connections (see **Supp. Fig. 6**). As the number of matrices increases, the distribution of A-B connection tends towards a normal distribution. We therefore fit the permuted A-B connection frequencies to a normal distribution with the MASS∷fitdist package in R-3.3 to obtain confidence intervals at +/- 1.96 standard deviations from the mean.

One possible explanation for signal enrichment of fusions between two tracks (or different elements within one track) is that the genomic distance between these elements is similar to the observed span distribution of the rearrangements. To test whether this was the case, we randomized the locations of the SINE elements (SINE-SINE fusions being the most enriched signal) and applied Swap using the randomized SINE track. We further ran Swap using only inter-chromosomal translocations, which removes any possibility of confounding by the rearrangement span distribution. We observed no enrichment or depletion for connections between the randomized SINE elements, using either the full model or the inter-chromosomal-only model (**Supp. Fig. 7d**). The degree of enrichment for SINE-SINE elements was nearly identical between the full model (1.19 fold-enrichment) and the inter-chromosomal only model (1.23 fold-enrichment).

### Break-invasion and double-break join models

We considered two simple background models based on the span distribution and the frequency with which each locus suffers rearrangements. The first model hypothesizes that the background probability is 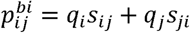, where *q*_i_ is the marginal probability of a rearrangement initiated in locus *i* and *s*_*ij*_ is the conditional probability that a break at *i* will connect to site *j*. Since we cannot distinguish between the start and end sites, we also add the reciprocal term, to yield a probability proportional to the local rate of retreatments connecting sites *i* and *j*. The marginal of the start site, *q*_i_, is determined from the empirical breakpoint density, *R*_i_, by applying preconditioned conjugate gradient descent optimization to the following problem:

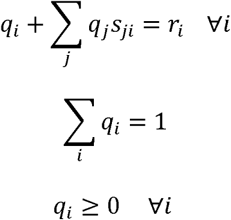

The conditional probability matrix is determined from the empirical span distribution, illustrated in **Fig. 2a**. The second model hypothesizes that the background probability is 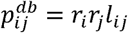, where *r*_*i*_ and *r*_*j*_ are the breakpoint densities and *l*_ij_ is a span factor connecting sites *i* and *j* found by solving the following constrained nonlinear optimization problem:

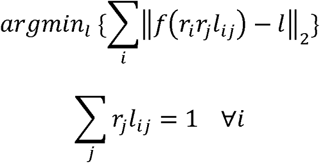

The function *ƒ* transforms the probability matrix to a span distribution function corresponding to the empirical distribution, *l* (**Fig. 2a**).

Both of these models have physical interpretations: in the first case, a genomic locus undergoes a break followed by invasion into another locus; and in the second case, both genomic loci undergo breaks followed by an erroneous join. We therefore termed the first model “break-invasion” and the second model “double-break join” (**Ext. Fig. 4a**). These physical interpretations are reminiscent of known DNA repair mechanisms. For example, non-allelic homologous recombination (NAHR)^17^ involves strand invasion after an initial break, similar to the physical interpretation of the break-invasion model. Conversely, non-homologous end joining (NHEJ) and MMEJ^18^ involve two or more breaks that are fused, similar to the physical interpretation of the ‘double-break join’ model.

Each of these models implies different distributions of fusions across the 2D map, enabling us to determine which model best fit the fusion patterns we observed. For example, simple, short rearrangements fit the probability distribution described by the break-invasion model, whereas complex interchromosomal rearrangements fit the distribution described by the double-break join model (**Ext. Fig. 4b**).

### Assessing the significance of somatic rearrangement fusions

To construct the probability matrices of the break-invasion, 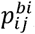, and double-break join, 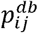,models, we divided the genome into bins containing a target of 100 rearrangements per bin. To avoid cases in which a cluster of rearrangements is divided into two bins, we imposed a minimal distance between breakpoints of 2 Kbp; if a bin boundary falls between two breakpoints not meeting this condition the bin is extended until the condition is met. The distribution of bins sizes is shown in **Supp. Fig. 8a;** the median bin size is 467 Kbp. Correspondingly, the distribution in the number of breakpoints per bin is shown in **Supp. Fig. 8b**; the median number is 91. The normalized distribution of number of breakpoint is the parameter *r*_*i*_ used to construct the two models. After binning the genome, we constructed the rearrangement matrix, *k*_*ij*_, by assigning each rearrangement in our dataset to a tile. Each sample was only allowed to contribute up to one rearrangement per tile.

The overall background rate of events is represented by a linear combination of the break-invasion and double-break join models. We defined the local rearrangement probability as 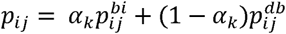, where the linear combination is taken over a set of parameters *α*^*k*^. We chose to use the distance between breakpoints as a natural choice for the classifier in this two-dimensional genomic representation. We divided the 2D space into short (≤1 Mbp), long (>1 Mbp), and inter-chromosomal translocations, and obtained the values of *α*^*k*^ by minimizing the Bayesian Inference Criteria (BIC). A list of recurrent rearrangements was then generated by calculating a p-value in each tile with a binomial test statistics against *k*_*ij*_, followed by control of multiple hypotheses using the Benjamini-Hochberg false discovery rate procedure.

One possibility is that these recurrent juxtapositions were mechanistically rather than selectively favored, for instance because they have high rates of microhomology, enabling microhomology-mediated repair. In fact, the opposite is true: only 14% of the recurrent fusions exhibited more than 2 bases of microhomology, significantly lower than the genome-wide average (25%; p<10^-4^).

### Power calculations

To analyze the number of tumor-normal pairs needed to reach saturation in the detection of fusions we employed a binomial power model^63^. We defined a null distribution, H_null_ ~ Binomial(*N,p*_null_), where *p*_null_ = 1-(1- *p*_90_)^*m*^ is the probability of a patient having at least one rearrangement, *p*_90_, is the 90^th^ percentile value of *p*_ij_ from our background model probabilities, and *m* is the median number of rearrangements per sample. The two-dimensional genomic fusions map was divided into 100 x 100 kbp tiles in this power analysis.

We performed the analysis first as a function of the distance between breakpoints with median number of rearrangements per sample of the entire cohort (**Fig. 6a**). The second analysis was performed as a function of the median number of rearrangements per sample, spanning values represented by the ICGC histologies with more than 15 samples (**Fig. 6b**). For each total number of tumor-normal pairs, *N*, the general procedure involved: 1) finding the minimal number of patients needed to reach significance level of *p* < 0.1/(# of tiles) based on H_null_; 2) using this value, calculating the minimal rate above background, *r*, that yields 90% power of the alternative distribution, H_alt_ ~ Binomial(*N,p*_null_ + *r*); 3) calculating contour lines of constant value rates above background.

### CESAM analysis

CESAM integrates rearrangement-derived breakpoints with RNA-seq data (FPKM-UQ) to identify expression changes associated with breakpoints in cis, as previously described ^4^. In brief, normalized RNA-seq expression is regressed on a rearrangement breakpoint matrix, using tissue-type, total number of rearrangements and principal components of the breakpoint matrix as covariates. Expression data was dosage-adjusted prior to the analysis by normalizing the expression level of each gene (FPKM) to the copy number level of the gene in each tumor sample. This was done to remove effects due to copy-number dosage effects, i.e. not attributable to *cis*-effects.

Three types of CESAM analyses were pursued to identify recurrent breakpoints associated with expression changes in cis: i) genomic regions of clustered breakpoints (termed ‘breakpoint_cluster’ in the CESAM analysis), ii) breakpoint fusion regions (termed ‘fusion_analysis’), and iii) TAD-bound breakpoints separated by germ-layer (termed TAD-bound’). For each analysis, only regions with at least three tumors having associated RNA-seq data. The ‘breakpoint_cluster’ and ‘fusion_analysis’ was performed on the complete pan-cancer set and for each histology type separately (annotated e.g. “PCAWG” and “1D_ORGAN_Lymph”, respectively in **Supp. Table 3** and **6**).

The association between adjusted gene expression changes of CESAM hits with breakpoints was assessed by computing the average of copy number-adjusted gene expression fold-change for each histology type.

To assess whether CESAM hits were associated with juxtaposition of normally distant enhancer elements, the distant breakpoint of a rearrangement (here defined as the breakpoint most distant to the gene-of-interest) was intersected with tissue-matched enhancer regions ^64^ with a window of +/- 20 Kbp. Significance was assessed by random shuffling of breakpoint positions on the mappable genome (alpha<0.1) and annotated as “cis_activating_enhancer” (**Supp. Table 3**).

### Rearrangement-types and effect on expression, enhancer-distance and TSGs

For each cluster of rearrangements *c*_*i*_, the genomic centroid position of the breakpoints was taken and the most deregulated gene within a window of +/- 1 Mbp of the centroid was identified.

To calculate the fold-change expression, for each of ci, with a set of tumor samples having a breakpoint at the cluster (denoted SV+) and a set of control samples without a breakpoint in the cluster (denoted rearrangement-), expression fold change was calculated as the median of the expression of SV+ samples divided by the median of the expression of SV-samples. A randomized background set was calculated for each ci by random sampling (n=100) a breakpoint from the PCAWG rearrangement set and computing fold-change as above with the same set of SV+ and SV-samples.

The distance to the nearest tissue-specific enhancer was computed as described under CESAM analysis.

Biallelic inactivation was assessed as described in detail in our accompanying PCAWG paper (Radhakrishnan et al) by requiring a copy-number loss of one allele and a disruptive rearrangement of the other allele. Only curated tumor-suppressor genes were assessed, as described in Radhakrishnan et al. Enrichment of biallelic inactivation for each rearrangement cluster type was assessed by comparing the frequencies to a permuted set (Fisher’s exact test, n=1,000), showing enrichment of biallelic inactivation at DEL-type (p<0.005), NEUTRAL-type (p<0.001) and FRAG-type (p<10^-13^) and depletion of AMP-type (p<10^-10^) and FUSION-type (p<10^-26^) rearrangement clusters.

### Rearrangement span and effect on TADs

The association between rearrangement size, TAD-disruption, and gene expression was assessed for TAD-bound CESAM analysis rearrangements. Each rearrangement was associated with the gene expression of the most significant CESAM-identified gene and separated into TAD-disrupting and TAD-preserving. Fold-change expression change was associated with rearrangement span (50 Kbp sliding bins with 25 Kbp overlap) with a 2nd order polynomial fit.

### Statistical calculations and software

Statistical calculations for performed using R-3.3.2. Student’s t-test calculations were obtained from stats∷t.test. Wilcox rank-sum tests were obtained from stats∷wilcox.test. The Spearman rank correlation coefficients were calculated using stats∷cor. The negative binomial and Poisson distributions were fit to the breakpoint count histogram using MASS∷fitdistr.

